# Differential DNA methylation and transcriptome revealed enhanced photosynthetic recovery and ROS modulation to promote cold tolerance in *Oryza sativa* L. ssp. *indica*

**DOI:** 10.64898/2026.04.30.721858

**Authors:** Vishal Roy, Rukshar Parveen, Pratiti Dasgupta, Shubho Chaudhuri

## Abstract

*Indica* rice, being a tropical crop, is highly sensitive to cold temperatures. Cold stress affects vegetative growth, photosynthetic efficiency and reproductive features. Screening diverse landraces to identify novel alleles associated with cold tolerance can be an approach to develop cold resilient indica rice. We have characterized a boro germplasm, CB1, with an efficient germination rate and growth vigor when treated with chilling temperatures. CB1 seedlings show a higher survival rate compared to IR36 when subjected to prolonged chilling stress. Biochemical analysis indicated efficient ROS modulation, higher chlorophyll content, enhanced photosystem II efficiency and unique stomatal traits, leading to higher relative water content in CB1 plants during stress and recovery. Germplasm-specific DNA methylome and transcriptome uniquely regulate the expression of chlorophyll biosynthesis, photosystem, & light harvesting complex and ROS scavenger genes in CB1 seedlings during stress recovery. Interestingly, high D1 protein turnover in CB1 promotes damage repair of PSII for efficient photosynthesis. Furthermore, key transcription factors for stomatal development and expression of photosynthetic genes were upregulated in CB1 during stress recovery. Notably, higher expression of *GLK1* and enrichment of GLK1 targets were observed in CB1 plants during chilling stress and recovery. Taken together, our results indicate that GLK1 and WRKY40 negatively regulates the inhibitory role of ABI5 for efficient photosynthetic recovery to enhance cold tolerance in CB1 rice.

## INTRODUCTION

Cold stress has been shown to have a global impact on rice production in high-altitude regions of tropical countries. Among the two Asian cultivated rice varieties, *indica* varieties are more sensitive to cold temperature compared to japonica when grown at higher elevated regions (Kumar et al., 2017). Boro rice grown in India during the dry winter season faces chilling stress (0°C-12°C) at the seedling stage in the North-eastern and western hill states of the Himalayas. Cold temperatures cause severe effects on the vegetative growth of rice plants, starting from rice germination, seedling growth and vigor, reduced tiller formation (El-Refaee et al., 2024). During the reproductive stage, cold stress affects flowering and panicle formation, that causes poor grain quality and loss of grain yield (Loitongbam et al., 2017). Thus, it is important to characterise different rice genotypes to identify rice germplasm having allelic variation that contributes to cold tolerance.

Although considerable research has been done in past to understand cold stress response, the mechanism of stress recovery has been less explored. The plant-stress-recovery mechanism is very crucial for recovering from damage caused by stress and reactivation of physiological processes for plant growth and development (Crisp. et al., 2016). During water stress, plants undergo extensive morphological, physiological, and biochemical changes, which, upon rehydration, recover its photosynthesis, stomatal conductance, growth-promoting hormone signalling, and gene expression (Xu et al., 2010). Similar results were obtained for *Zea mays,* where plants after being subjected to drought and cold stress in combination, displayed efficient recovery of photosynthetic parameters and stomatal conductance (Guo et al., 2021). Exogenous application of melatonin and brassinosteroids accelerates cold stress recovery in melon and cucumber, respectively, by reviving photosynthesis and activating antioxidant scavengers (Jiang et al., 2013; ZHANG, 2017).

Although being more susceptible to cold stress, studies have identified specific *indica* varieties and hybrids that showed varying degrees of tolerance to cold temperatures at different stages of plant development. Liyouyuchi, an *indica* hybrid rice, demonstrated superior cold tolerance, with reduced reactive oxygen species (ROS) accumulation, higher survival rate and higher grain yield during cold stress compared to its parental lines (Zhang et al., 2025). Similarly, another *indica* genotype, IRGA 959-1-2-2F-4-1-4-A, showed a tolerance phenotype with high photosynthetic performance after recovering from cold exposure (Adamski et al., 2016).

In plants, epigenetic alterations within a group can be a rich resource for genetic diversity and natural variation. Rice has rich germplasm resources with variations in morphological, agronomic, and adaptive traits. Significant variations in DNA methylation within rice germplasms can be a key epigenetic factor that regulates phenotypic traits (Kotaro Miura, 2009). Studies have shown that DNA methylation modulates chromatin accessibility for gene expression, transposon silencing for maintaining genomic integrity, response to environmental cues and stress memory. Recent studies have shown that abiotic and biotic stresses can cause changes in DNA methylation in plants. However, most of the stress-induced DNA methylation reverts to its initial state after removal of the stress signal. DNA methylation marks, which are retained after stress removal, can be inherited meiotically and contribute to multi-generational adaptation against environmental stress (Song et al., 2025; Wibowo et al., 2016). Therefore, studying DNA methylation dynamics in different rice germplasms is important to identify loci exhibiting specific methylation patterns that confer cold tolerance in *indica* rice.

In this study, we have characterised two *indica* rice germplasms, cold sensitive IR36 and cold tolerant CB1, using comparative whole genome transcriptome and DNA methylome to understand the stress recovery mechanism against chilling stress. The physiological, biochemical and transcriptome data support efficient photosynthetic efficiency, ROS management and stomatal dynamics in cold tolerant CB1 during stress recovery. Further, significant DNA methylation variations and distribution patterns were observed between the two rice germplasms when subjected to chilling stress or post stress recovery, to regulate gene expression. Collectively, our results suggested that unique genetic and epigenetic modifications govern chilling stress tolerance and adaptability in CB1 plants.

## RESULTS

### Phenotypic comparison of IR36 and CB1 indica rice under normal growth conditions

Initially, we compared the phenotypic characteristics of two indica rice germplasms, IR36 and CB1. Under normal growth conditions, CB1 plants were significantly taller (206±5 cm) than IR36 (130±10 cm) **(Fig. 1A, i–ii)**. CB1 plants showed approximately two more internodes compared to the typical 4–5 internodes in IR36 **(Fig. 1A, iii)**, with significantly elongated individual internodes **(Fig. 1A, iv)**. At the flowering stage, both the rice varieties displayed distinct panicle morphologies **(Fig. 1B, i)**. CB1 exhibited a 50% reduction in total panicle number relative to IR36 **(Fig. 1B, ii)**. Furthermore, panicle length was significantly longer in IR36 plants compared to CB1 **(Fig.1B, iii),** with a higher number of spikelets per panicle **(Fig.1B, iv)**.

**Figure 1:**
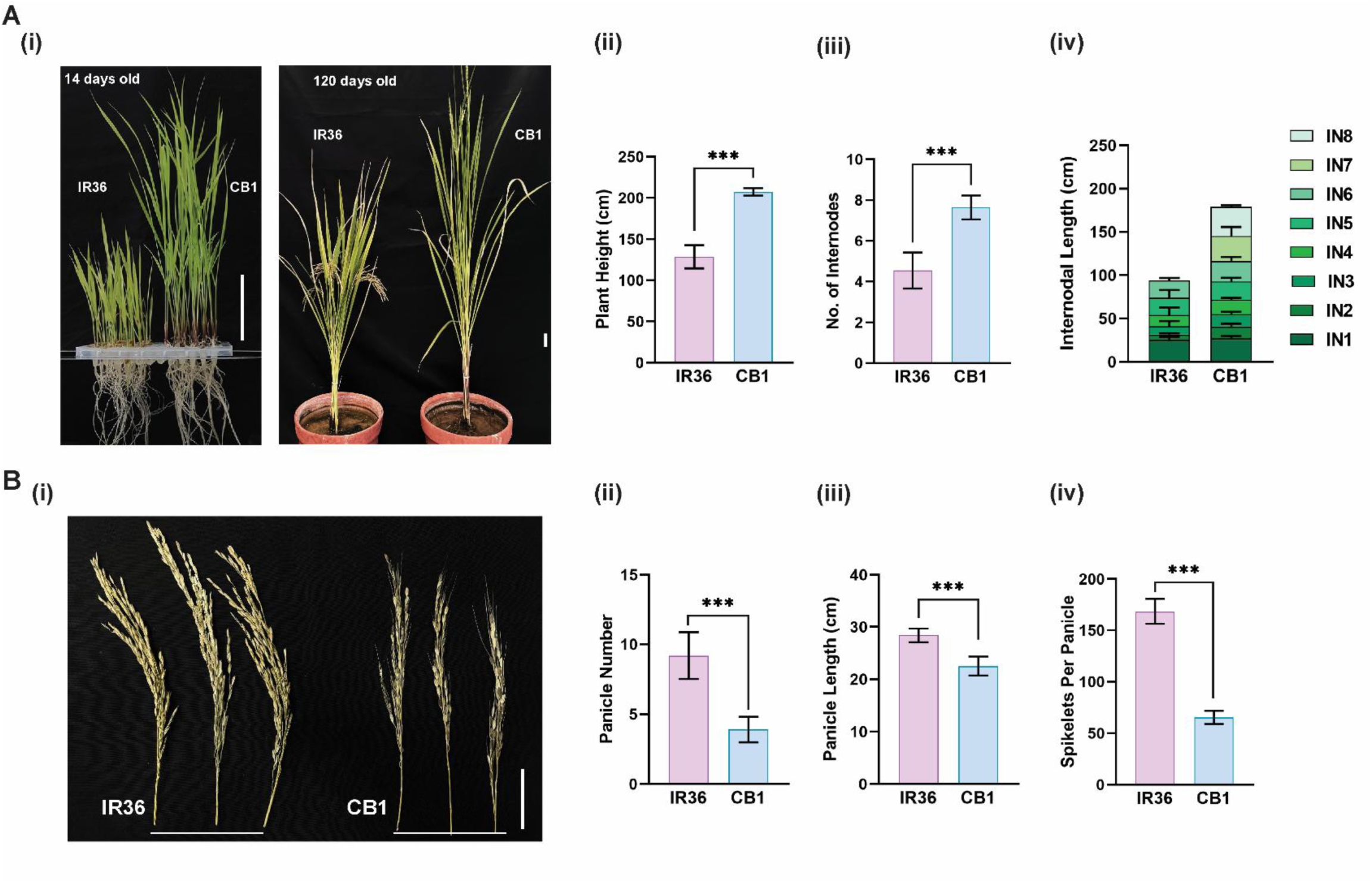
Morphological difference between IR36 and CB1 plants. A. **(i)** Pictorial representation of IR36 and CB1 rice at 14 days old and 120 days old. Scale bar = 5 cm. Bar graph showing **(ii)** the average height of IR36 and CB1 plants, **(iii)** number of internodes, **(iv)** internodal length**. B. (i)** Panicle morphology of IR36 and CB1. Bar diagram showing **(ii)** panicle numbers**, (iii)** panicle length**, (iv)** spikelets per panicle. The data represents an average of 20 matured plants. Error bar represents mean ± S.D. (*n* = 20). The significance of the data was measured by two-tailed paired t-test, the level of significance was represented by \*\*\**P <* 0.001.

### CB1 seedlings show higher survival with better water retention post-chilling stress

14 days old seedlings of IR36 and CB1 were subjected to chilling stress (8°C for 24, 48, and 72 hours) followed by 7 days of recovery at 28°C. While CB1 plants maintained a 90% survival rate after 7 days of recovery post-chilling stress, survival of IR36 seedlings declined significantly after cold exposure: less than 60% survived after 24 hours and 10% after 48 and 72 hours **(Fig. 2A-B)**. Furthermore, CB1 retains a higher relative water content (RWC) during recovery, maintaining 80% RWC after 72 hours stress, in contrast, RWC decreased significantly (20%) in IR36 **(Fig. 2C).**

**Figure 2:**
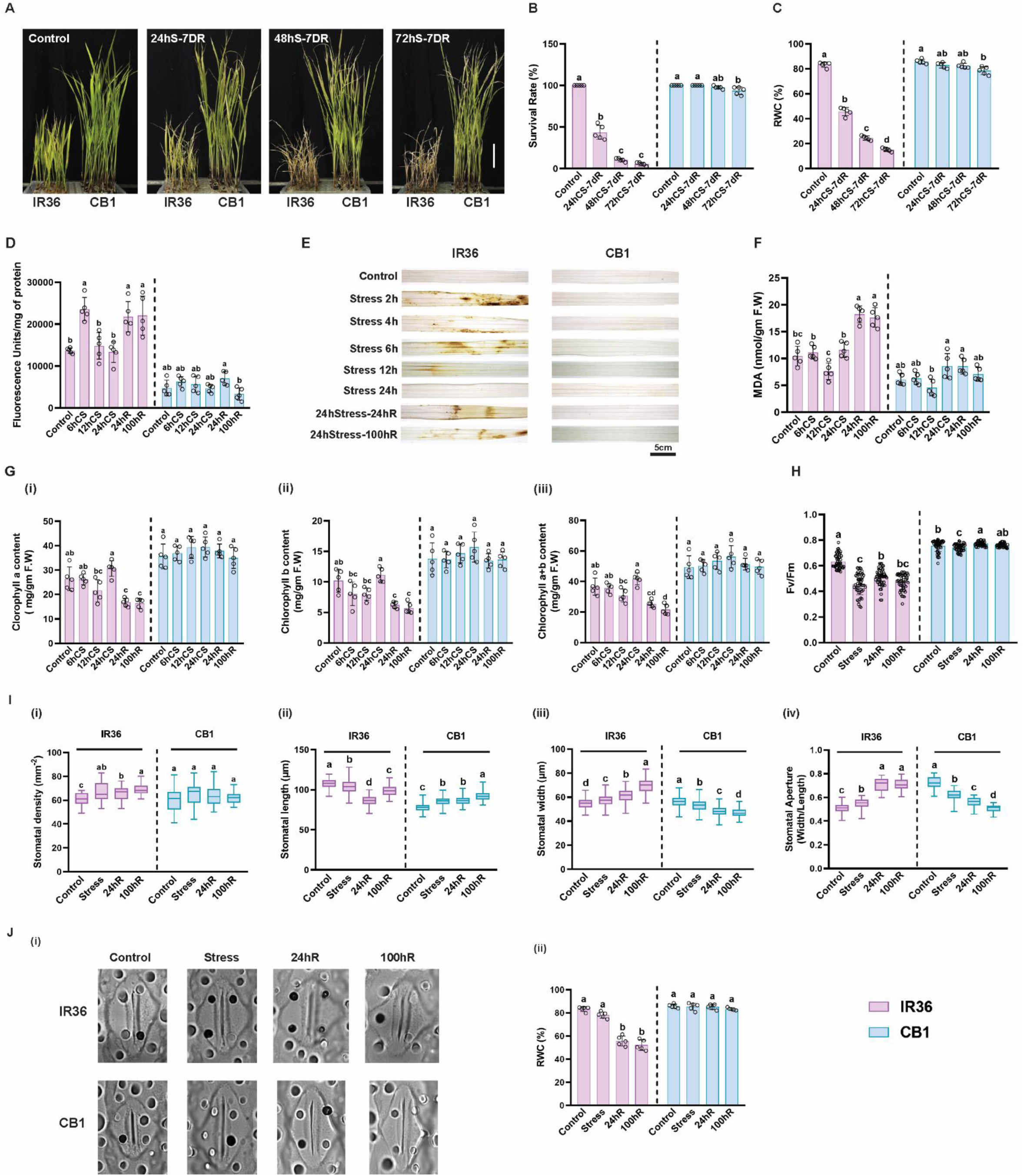
CB1 plants showed better survival against chilling temperature. **A.** Image showing 14 days old IR36 and CB1 subjected to chilling stress at 8°C for 24h, 48h and 72h followed by recovery for 7 days under control condition (28°C). Scale bar = 5 cm. **B.** Bar diagram for IR36 and CB1 plants showing survival rate (% survival), **C.** Bar graph showing relative water content (%) in IR36 and CB1, after 7 days of recovery post chilling stress for 24h, 48h, 72h. **D.** Quantitative analysis of total ROS content **E.** H_2_O_2_ accumulation on the leaves of IR36 and CB1, **F.** Total MDA content at different stages of chilling stress and recovery in 14 days old seedling of IR36 and CB1. **G.** Chlorophyll content (Chl A and Chl B) was estimated at different stages of chilling stress and recovery in 14 days old seedling of IR36 and CB1 **(i)** Chlorophyll A, **(ii)** Chlorophyll B and **(iii)** Chlorophyll A+B. **H.** Maximum quantum yield of photosystem II (Fv/Fm) estimation at control, 24h chilling stress and recovery phase (24h and 100h) in 14 days old seedling of IR36 and CB1. Stomatal traits were measured on fully expanded 3^rd^ leaf of IR36 and CB1 during control, chilling stress (24h) and recovery (24h and 100h). **I. (i)** Stomatal density, **(ii)** stomatal length, **(iii)** stomatal width and **(iv)** stomatal aperture. **J. (i)** Representative confocal image of individual stomata of IR36 and CB1 of control, chilling stress and recovery (24h, 100h). Scale bar =10 µm **(ii)** Relative water content (%) of IR36 and CB1 seedlings during control, chilling stress (24h) and recovery phase (24h, 100h). For each bar, the error bars represent mean ± S.D, except for stomatal parameter where bar represents median of the data from five independent experiments (*n* = 5). The significance of the data was measured by one-way ANOVA with Tukey’s post hoc test.

These results indicated better performance of CB1 plants in maintaining water balance post chilling stress compared to IR36.

### CB1 exhibits enhanced ROS scavenging and maintains photosynthetic efficiency during chilling stress and recovery

Chilling stress causes increased ROS accumulation in plants, resulting in oxidative damage (Sharma et al., 2012). In IR36 plants, total ROS increased significantly after 6 hours of chilling stress, declined at 12 and 24 hours, followed by a further increase during post-stress recovery (24 hours and 100 hours). Unlike IR36, CB1 seedlings showed better ROS scavenging with no significant change in the total ROS content under chilling stress or recovery phase **(Fig. 2D-E)**. Furthermore, MDA content was increased in IR36 during recovery phase, whereas no change in MDA content was observed in CB1 plants during stress or recovery **(Fig. 2F)**. Collectively, these results indicated that the CB1 plants have highly active ROS scavenging machinery to protect cellular components from oxidative damage during chilling stress.

IR36 leaves showed visible chlorosis after chilling stress compared to CB1. During chilling stress and recovery, no change in chl *a*, chl b and total chlorophyll (a+b) contents was observed in CB1 seedlings. However, chlorophyll content decreased significantly during recovery in IR36. **(Fig. 2G, i-iii)**. Further, maximum quantum yield of photosystem II (Fv/Fm ratio) decreased significantly in IR36, whereas CB1 seedlings showed minimal effect under all conditions. These results indicated compromised photosynthetic efficiency in IR36 after chilling stress compared to CB1 seedlings **(Fig. 2H)**.

Additionally, stomatal traits differed distinctly between the two rice germplasms under stress and recovery. Interestingly, stomatal density increased in IR36 plants during stress and recovery; however, no significant change was observed in CB1 plants **(Fig. 2I, i)**. Notably, stomatal length decreased **(Fig. 2I, ii),** width increased **(Fig. 2I, iii),** causing an increase in stomatal aperture (**Fig. 2I, iv & Fig. 2J, i**) in IR36 during the stress and recovery phase. However, CB1 plants showed a contrasting trend with increased length and a significant decrease in stomatal width, resulting in closed or partially open stomata to maintain leaf RWC (**Fig. 2J, ii**).

### Comparative transcriptome between IR36 and CB1 under chilling stress and recovery

To evaluate transcriptional changes, RNA-seq was performed using 14 days old IR36 and CB1 seedlings subjected to 24 hours of chilling stress and 24 hours of recovery. During chilling stress, 11915 DEGs (7201 up and 4714 down-regulated) and 11107 DEGs (6748 up and 4359 down-regulated) were identified in IR36 and CB1 respectively, with 7979 DEGs common between the two germplasms. During recovery, 9393 DEGs (3791 up and 5602 down-regulated) and 11291 DEGs (4995 up and 6296 down-regulated) were identified in IR36 and CB1, respectively of which 6530 were common (**Fig. S1.A-F**).

Gene Ontology (GO) analysis of Upregulated common DEGs revealed enrichment of terms related to “stress responses’’ “response to temperature stimulus” and “hormone signalling”. Downregulated DEGs were enriched for “plastid organization”, “photosynthesis light harvesting photosystem I”, and “chloroplast organization”. For IR36 unique upregulated DEGs, GO enrichment was for “response to cold”, “oxylipin biosynthetic process”, whereas “triterpenoid and terpenoid biosynthesis”, “Photosynthesis light harvesting” for downregulated DEGs. Unique upregulated DEGs for CB1 were enriched in “response to auxin”, “response to brassinosteroid”, whereas downregulated DEGs were enriched for “photosynthesis”, “macromolecule methylation”, “nitrogen compound catabolic process”, and “membrane lipid metabolic process”. During recovery, upregulated common DEGs were enriched for “plastid organization” and “photosynthesis”, whereas downregulated DEGs were categorized under “response to JA”, “response to ethylene” and “hormone signaling”. Upregulated Unique DEGs in IR36 were enriched in “terpenoid metabolism”, “photosynthesis light reaction” and “isoprenoid biosynthesis” whereas downregulated DEGs were involved in “response to abscisic acid”, “reproductive structure development” and “gibberellin catabolic process”. Similarly, for CB1 upregulated DEGs were enriched in “pigment biosynthesis”, “photosynthesis light harvesting photosystem I”, “photosynthesis light harvesting”, “chloroplast organization”, “chlorophyll biosynthesis” and “cell wall organization”. GO enrichment for downregulated DEGs of CB1 are “response to salicylic acid”, “response to cold”, “oxygen-containing compounds” and “brassinosteroid signalling” **(Fig. S1.G).**

### Genome-Wide DNA methylation dynamics in IR36 and CB1 plants

Whole genome analysis revealed that ^m^CG and ^m^CHG were predominantly enriched in centromeric and chromosome flanking region in both varieties (**Fig. 3A**). Interestingly, a significant difference in methylation level was observed when compared between the two rice germplasms **(Fig. 3B)**. Under control conditions, CB1 plants have lower ^m^CG (37% vs. 40.6%), but higher ^m^CHH (38.4% vs 33.8%) compared to IR36. ^m^CG declines (37% to 35.7%) during chilling stress in CB1, whereas ^m^CHH increases from 38.4%. to 40.2%. Chilling stress also increases ^m^CHH in IR36 from 33.8% to 35.9%. Upon stress recovery, CB1 fully restored baseline methylation levels, whereas IR36 plants showed reduced mCG and elevated mCHH with respect to baseline **(Fig. 3B)**. Region-specific profiling showed that IR36 maintained higher methylation levels in upstream, gene body, and downstream regions under control and stress conditions relative to CB1 **(Fig. 3C, i-iii)**. However, methylation significantly decreased in these regions during recovery in IR36. Gene body showed preferentially higher ^m^CG methylation than ^m^CHG and ^m^CHH, with higher ^m^CG in IR36 than CB1. These analyses suggested the presence of germplasm-specific DNA methylation dynamics for stress response and recovery in IR36 and CB1.

**Figure 3:**
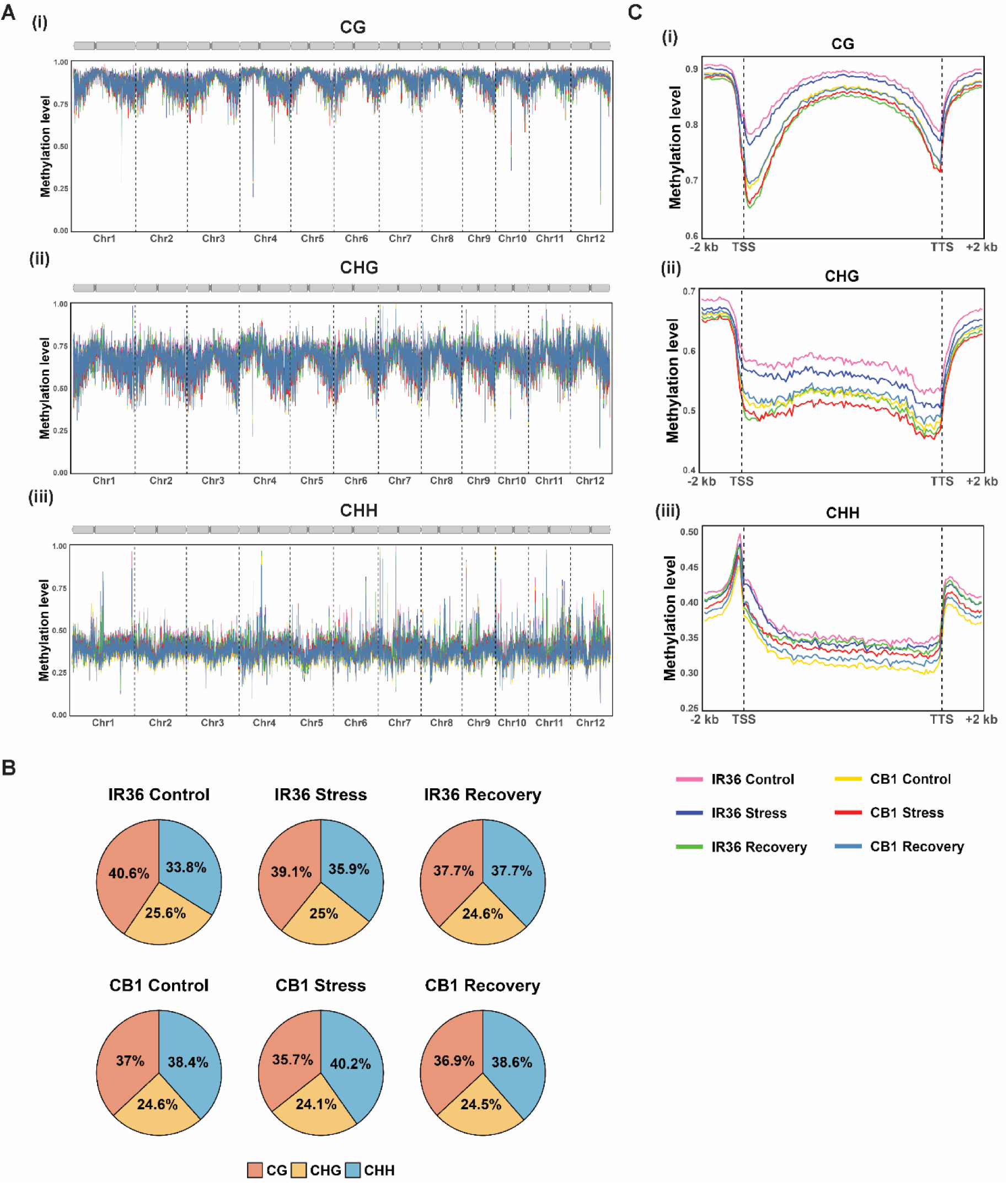
Genome wide DNA methylation in IR36 and CB1. **A.** Metagene plots showing chromosomal distribution of average methylation signals across 12 chromosomes, calculated in 50kb sliding windows for **(i)** ^m^CG **(ii)** ^m^CHG and **(iii)** ^m^CHH contexts under control, stress and recovery. **B.** percentage of methylation in three sequence contexts (CG, CHG, CHH) under control, stress and recovery. **C.** Changes in methylation level in 2 kb upstream, gene body and 2kb downstream region of genes in all three-sequence context during control, stress and recovery condition.

### Integrated transcriptome and DNA methylome revealed enrichment of photosynthesis and chloroplast organization in CB1

Initial investigation of differentially methylated regions (DMRs) between the two germplasms when grown under normal growth condition, revealed distinct difference in DMRs profiling with CB1 plants having large number of DMRs in all three-sequence context (**Fig 4A, i**). CG context DMRs (hyper or hypomethylated) were found associated with large number of DEGs (up-or down regulated) in CB1 plants compared to IR36, followed by CHG and CHH context (**Fig 4A, ii**). DMRs with all three methylation marks showed differential distribution in promoter, gene body and intergenic region (**Fig 4A, iii**). Although CHG or CHH-DMRs showed unbiased distribution across promoter, gene body and intergenic region, with CHG-DMR mainly enriched in the intergenic region.

**Figure 4.**
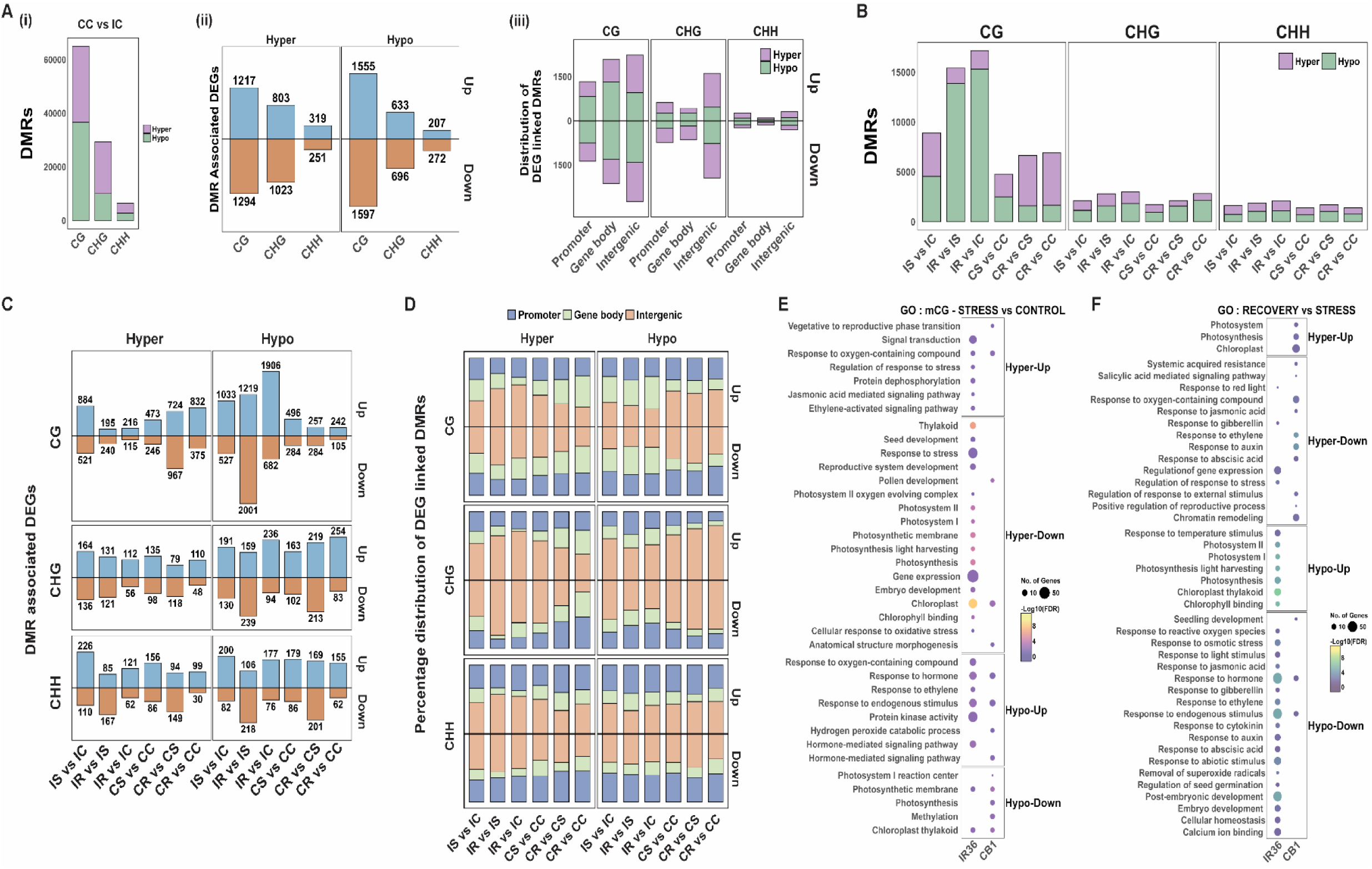
DMR analysis and its association with DEGs during stress and recovery in IR36 and CB1. **A.** Differentially methylated regions were analysed between two germplasms, CB1 vs IR36 under control condition. **(i)** Hyper and Hypo-methylated DMRs in three sequence contexts**, (ii)** DMR-associated DEGs, **(iii)** Distribution of DMRs in different genomic regions. **B.** Hyper and Hypo-methylated DMRs in three sequence contexts under Stress vs Control, Recovery vs Stress and Recovery vs Control conditions in IR36 and CB1plants. **C.** DMR-associated DEGs of three sequence contexts in CB1 and IR36 under Stress vs Control, Recovery vs Stress and Recovery vs Control conditions. **D.** Distribution of DMRs in different genomic regions of DEGs under Stress vs Control, Recovery vs Stress and Recovery vs Control conditions. **E-F.** Dot plot showing Gene Ontology enrichment analysis of DMRs associated DEGs in IR36 and CB1 under stress and recovery. Increasing circle size is positively correlated with the number of genes under respective GO terms while colour bar corresponds with -Log_10_FDR values.

Next, DMRs were identified for stress vs. control; recovery vs. stress and recovery vs control for individual plants. Most identified DMRs were predominantly associated with the CG sequence context. In IR36, 12,674 DMRs were identified during chilling stress, however low number of DMRs were identified for CB1. Interestingly, majority of the DMRs in IR36 were hypomethylated in CG context during recovery (**Fig 4B**). However, CB1 plants have fewer DMRs, mostly hypermethylated at CG. We have also analysed change in methylation status during recovery compared to the baseline methylation (control). With respect to control condition, IR36 stress recovered plants retained hypomethylated-DMRs, whereas for CB1, majority of DMRs after recovery were hypermethylated.

To link methylation dynamics with gene regulation during chilling stress and recovery, DMRs were mapped to DEGs in IR36 and CB1. In IR36, during chilling stress and recovery, a large number of upregulated and downregulated DEGs were identified, predominating associated with hypomethylated DMRs in CG context. In contrast, less number of DMRs, predominantly hypermethylated at CG, were mapped to DEGs in CB1 during chilling stress or during recovery. Comparison between recovery and control has also identified significant number of upregulated DEGs enriched with hypomethylated DMRs in IR36 and hypermethylated DMRs in CB1. Compared to CG context, very less number of CHG and CHH context DMRs were associated with DEGs under all conditions in both the plants (**Fig. 4C**). Furthermore, region specific distribution of DMRs across within DEGs, changes during chilling stress and recovery. In IR36, most of the hypermethylated DMRs are distributed in intergenic region during chilling stress and hypomethylated DMRs in the gene body during recovery. In CB1, hypomethylated and hypermethylated DMRs found majorly at intergenic regions under all conditions **(Fig. 4D)**.

Transposon element (TE) silencing is highly regulated through DNA methylation. TEs associated with genic regions remain highly methylated in all three-sequence context during control condition in IR36, that significantly decreased during chilling stress and recovery. In contrast, TEs in genic region are less methylated in CB1 plants and methylation at CG and CHG significantly reduced, while CHH increased during chilling stress. All three methylation marks revert back to baseline during recovery in CB1 **(Fig. S2.A)**. Similarly, for nongenic-TEs, methylation in all three contexts found highest in IR36 under control and maintained during stress or recovery. For CB1, only ^m^CHH level increases during stress and retuned to baseline in recovery **(Fig. S2.A)**. Further, more than 50% of CG context and 70% of CHG and CHH context DMRs were associated with TEs in all conditions **(Fig. S2.B)**. Among the TE associated DMRs, hypomethylated CG context DMRs predominate compared to CHG or CHH context, in most of the cases **(Fig. S2.C)**. Most TE-overlapping DMRs were mainly enriched at intergenic regions **(Fig. S2. D)**.

GO enrichment for upregulated DEGs having hyper-DMRs showed enrichment of terms like “response to stress”, “JA and ethylene signaling” in IR36 and “vegetative-to-reproductive phase transition” in CB1 during chilling stress. Similarly, hyper-DMRs associated downregulated DEGs showed enrichment of terms like “thylakoid”, “photosystems, light harvesting”, “chloroplast”, “response to oxidative stress” in IR36, and “anatomical structure morphogenesis” and “chloroplast” in CB1. In IR36, upregulated genes with hypo-DMRs have enriched GO terms for “response to oxygen containing compound”, “hormone signaling pathway” and “protein kinase” during chilling stress, whereas “chloroplast thylakoid” and “photosynthesis membrane” for downregulated DEGs. Similarly, for CB1, “hydrogen peroxide catabolic process”, “response to endogenous stimulus” and “hormone mediated signaling” were associated with upregulated DEGs having hypo-DMR and “photosystem I reaction center”, “photosynthesis” and “methylation” for hypo-DMR linked downregulated DEGs **(Fig. 4E)**.

During recovery in CB1, upregulated DEGs linked to hyper-DMRs have enriched terms like “photosystem”, “chloroplast” and “photosynthesis”, and “hormone signalling”, “chromatin remodelling”, “response to oxygen-containing compounds”, and “reproductive processes” for downregulated DEGs. In IR36, hyper-DMRs linked with downregulated DEGs were enriched for “stress response”, “gibberellin response”. Similarly, hypo-DMR associated with upregulated genes in IR36 were enriched for “photosynthesis”, “LHCs”, “photosystems I and II”, “chlorophyll binding”, and “temperature response” and downregulated genes for “response to oxygen-containing compounds”, “hormone signalling”, “abiotic stress”, “ion homeostasis”, “calcium binding”, “post-embryonic development”, “seed germination”, “ROS removal” **(Fig. 4F)**. Integrated transcriptome and methylome analysis reveal that many essential cellular processes were differentially regulated by germplasm specific DNA methylation in response to chilling stress and recovery.

### CB1 plants efficiently modulate the expression of photosynthetic genes during chilling stress and recovery

DMR-DEG analysis revealed significant enrichment of chlorophyll metabolism, photosynthesis, photosystems (PS), and light-harvesting complexes (LHCs) under stress and recovery in IR36 and CB1.

#### Chlorophyll metabolism

Transcriptome and methylome analyses indicated that among four major differentially expressed chlorophyll biosynthesis genes, only *PORB* has CG-context DMRs within the promoter region and gene body. *PORB* expression decreased during chilling stress in both germplasms **(Table S1)**; however, a significant increase was observed in CB1 during recovery **(Fig. 5A, i-ii)**. During chilling stress, ^m^CG, ^m^CHH marks increased at the promoter region of *PORB* and ^m^CHG, ^m^CHH within the gene body in CB1 plants. In contrast, significant changes in ^m^CHG in the *PORB* gene body were observed for IR36. Upon recovery, both promoter and gene body methylation levels returned to their baseline in CB1, while IR36 maintained elevated methylation of CG and CHG in the gene body **(Fig. 5A, iii & S4.A)**.

**Figure 5.**
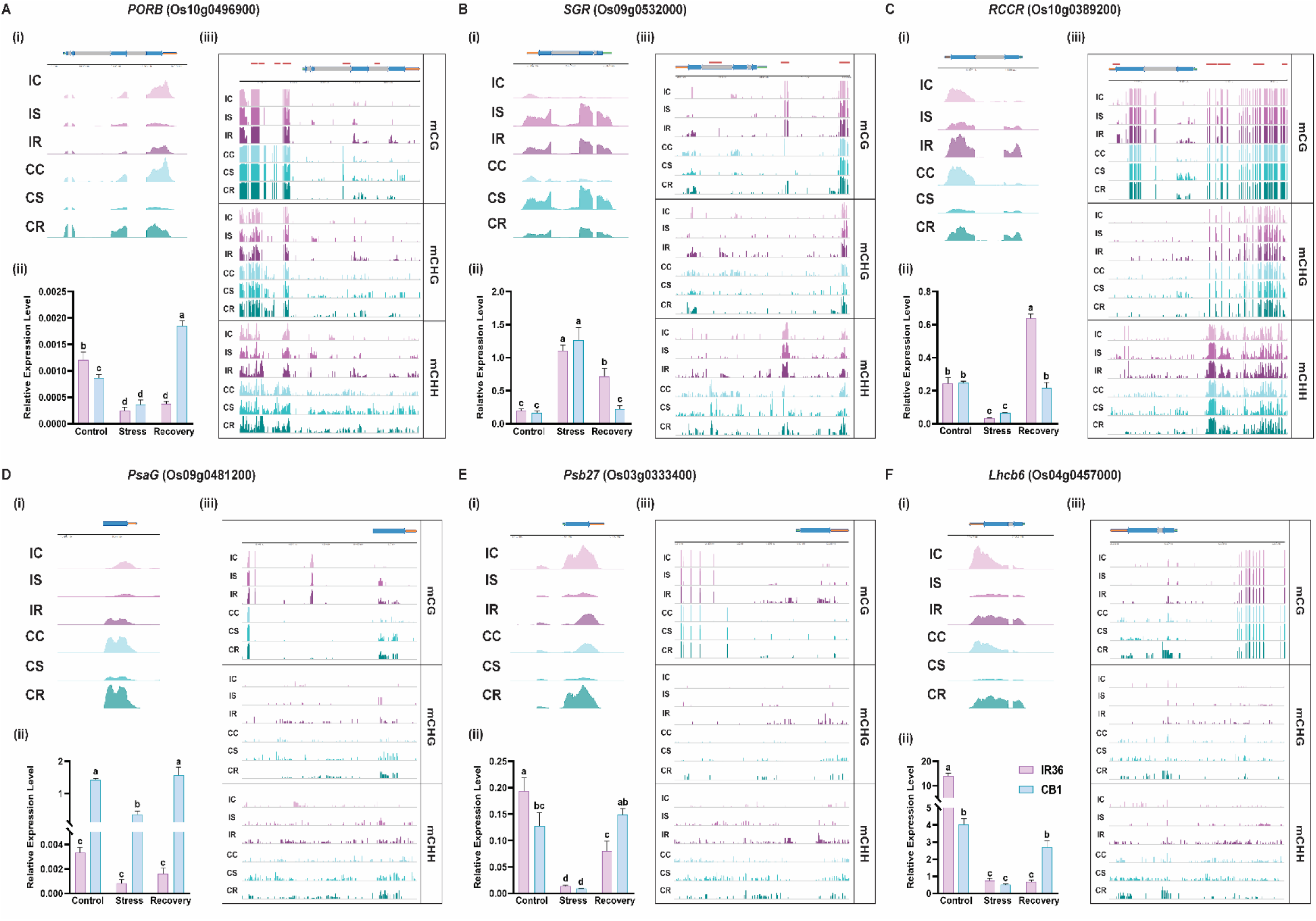
Changes in methylation and expression of photosynthesis related genes during chilling stress and recovery in IR36 and CB1. A-F. (**i**) RNA-seq coverage of respective genes in IR36 and CB1. (**ii**) RT-qPCR validation of expression of respective genes under control, stress and recovery. The error bars represent mean ± S.D. from three independent experiments (*n* = 3). The significance of the data was measured by two-way ANOVA with Tukey’s post hoc test. (**iii**) DNA methylation (CG, CHG and CHH) profiles within 2kb promoter and gene body under different conditions. IC: Control, IS: Stress, IR: Recovery for IR36 and CC: Control, CS: Stress, CR: Recovery for CB1. Red bars representing TE, Black Line (above) representing genome coordinates and horizontal bars represent gene body locations (Green: 5’UTR, Grey: Introns, Blue: Exons, Orange: 3’UTR).

#### Chlorophyll metabolism

Expression of key chlorophyll degradation genes (*NYC1*, *SGR*, and *NYC3*) were upregulated, and *RCCR* expression decreased under stress in both plants **(Table S1)**. Interestingly, expression of *SGR* and *RCCR* remained significantly higher in IR36 during recovery **(Fig. 5B, i-ii & 5C, i-ii)**. For the *SGR* locus, ^m^CHH level increased at the promoter, and ^m^CG and ^m^CHH significantly decreased at the gene-body during stress in IR36. During recovery, all three marks increased at the gene-body in IR36. For CB1, ^m^CHG and ^m^CHH increased at the promoter only during recovery, whereas ^m^CHG and ^m^CHH levels decreased significantly at the gene-body during stress and recovery **(Fig. 5B, iii & S4.B)**. For *RCCR* locus, all three methylation marks significantly increased in the gene body during stress in CB1 and reverted to baseline during recovery. In contrast, IR36 maintained significantly increased CHG and CHH methylation during stress and recovery **(Fig. 5C, iii & S4.C)**.

#### Photosystem

Most PS I and PS II genes identified in our DMR-DEG analysis were significantly downregulated under chilling stress in both germplasms **(Table S1)**. During recovery, expression was revived, reaching the highest levels in CB1 **(5D, i-ii; 5E, i-ii; S3.A, i-ii; S3.B, i-ii)**. Methylation at *PsaG* and *PsaL* loci orchestrated differently under stress and recovery when compared between two plants. At the *PsaG* promoter, IR36 showed no significant changes under stress but significant increase in ^m^CHG and ^m^CHH during recovery, whereas CB1 showed significant increase in the ^m^CHG and ^m^CHH both under stress and recovery. At *PsaG* gene body, ^m^CG and ^m^CHG level increased during stress in both germplasms. During recovery, ^m^CG and ^m^CHH levels were higher than baseline in IR36 whereas all three methylation marks increased progressively in CB1 **(Fig. 5D, iii & S4.D).** Similar differential methylation changes of CG, CHG and CHH were also observed for *PsaL* during stress and recovery in IR36 and CB1 plants **(Fig. S3.A, i & S4.E)**. The expression of PSII genes, *PsbR1 and Psb27*, were also enriched with different methylation marks during chilling stress and recovery. For *Psb27* locus, ^m^CHG and ^m^CHH gradually increased at the promoter under stress and recovery in IR36 plants however, all three marks initially decreased in gene body during stress but increased subsequently during recovery. For CB1, no change in methylation was observed at the promoter under stress or recovery; whereas^m^CHG increased during stress in gene body and ^m^CHH decreased during recovery **(Fig. 5E, iii & S4.G)**. *PsbR1* expression during stress and recovery is also regulated by differential methylation marks **(Fig. S3.B, iii & S4.F)**

#### LHCs

Multiple LHC genes associated with PSI and PSII were downregulated during chilling stress; however, expression recovered post-stress **(Table S1)**, with CB1 exhibiting better recovery **(Fig. 5F, i-ii; S3.C, i-ii)**. DNA methylation profiles at *Lhca4* and *Lhcb6* loci revealed distinct responses between the two germplasms. In CB1 plants, *Lhca4* and *Lhcb6* promoter methylation remained unchanged during chilling stress and recovery, except ^m^CHH level decreased at *Lhcb6* during recovery. In contrast, ^m^CHG and ^m^CHH increased at *Lhcb6* promoter during stress and maintained elevated during recovery in IR36, while ^m^CHG and ^m^CHH increased at *Lhca4* only during recovery. Gene body methylation (^m^CG and ^m^CHG) decreased significantly at *Lhca4* during stress in CB1and maintained below baseline in recovery, while these marks increased during recovery for *Lhcb6*. In IR36, all three methylation marks increased compared to baseline during recovery for both loci **(Fig. 5F, iii; S3.C, iii; S4.H-I)**.

### Higher expression of ROS scavengers in CB1 plants during chilling stress and recovery

Chilling stress–induced reactive oxygen species (ROS) often target nucleic acids, proteins, lipids, and chlorophylls (Sachdev *et al*., 2021). Plants adopt various mechanisms to scavenge ROS-induced toxicity, for maintaining redox homeostasis. Our biochemical studies revealed differences in ROS accumulation between two rice germplasms during stress and recovery. DMR-DEG analysis identified a milieu of antioxidant genes during stress and recovery. Based on their unique expression, these were categorised into four clusters (**Fig. 6A**, **Table S2**). Cluster I comprise class-III peroxidase genes that were upregulated under chilling stress in CB1 plants. Cluster II consists of catalase, ascorbate peroxidase, and peroxidase genes, showing maximum expression in IR36 under stress. Clusters III & IV included genes that were specifically upregulated during recovery while remaining downregulated under stress. Transcriptome data were validated using RT-qPCR on representative genes from Clusters I, II, and IV. The RT-qPCR results aligned with the transcriptome profiles, except for *Apx1*, showing significantly higher in CB1 during chilling stress compared to IR36 **(Fig. 6B, i-vi)**. Germplasm-specific DNA methylation changes at promoter and gene-body regions were observed during chilling stress and recovery (**Fig. S5.A-F**). Higher expression of antioxidant gene during stress and recovery suggests an efficient ROS scavenging mechanism in CB1 to protect oxidative damage.

**Figure 6.**
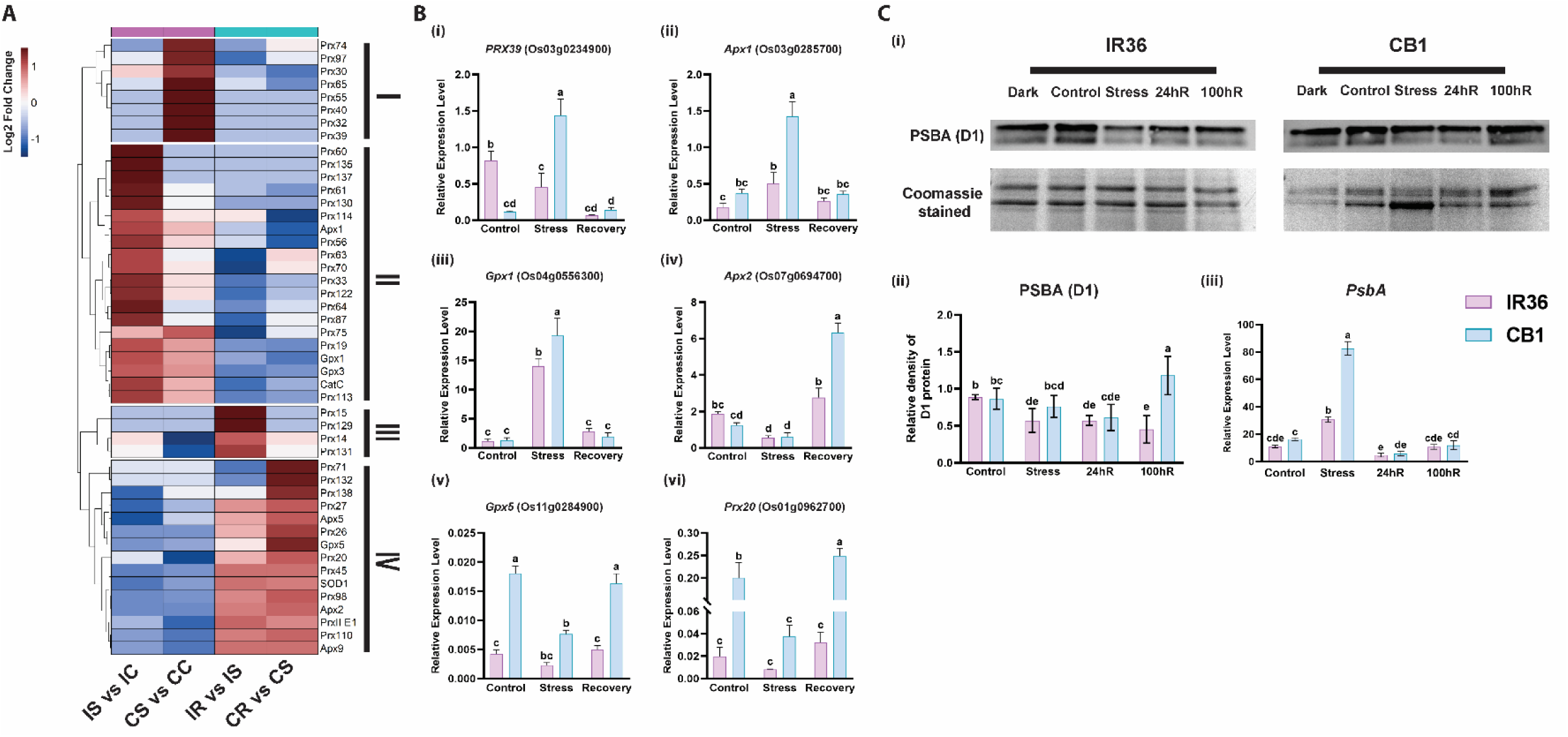
CB1 plants have efficient ROS scavenging mechanism. **A.** Heatmap of DMR associated differentially expressed antioxidant genes in IR36 and CB1 under chilling stress (IS vs IC and CS vs CC) and recovery (IR vs IS and CR vs CS). The cluster was categorized based on their unique expression patterns during stress and recovery period in IR36 and CB1. The heatmap was generated using log_2_ fold change values. **B. (i-vi).** RT-qPCR validation of the expression of some of the key antioxidant genes that are involved in scavenging stress induced ROS in chloroplast. The error bars represent mean ± S.D. from three independent experiments (*n* = 3). The significance of the data was measured by two-way ANOVA with Tukey’s post hoc test. **C. (i)** Western blot showing change in D1 protein level, **(ii)** quantitative analysis of band intensity of D1 protein, under control, chilling stress and recovery (24h and 100h). The data represents from three independent experiments (n=3) and **(iii)** RT-qPCR of *PsbA* (D1) gene. The error bar represents mean ± S.D. from three independent experiments (*n* = 3). The significance of the data was measured by two-way ANOVA with Tukey’s post hoc test.

Photoinhibition of PSII measures the rate of photodamage vs the rate of repair of PSII. Extreme environmental conditions cause oxidative damage to the PSII reaction center–binding protein D1, resulting PSII destabilization and reduction of photosynthetic efficiency. Western blot analysis indicated a decrease in D1 protein content during chilling stress in both the germplasms; however, CB1 fully recovered D1 protein content, whereas recovered IR36 plants maintained reduced D1 levels (**Fig. 6C, i–ii**). Furthermore, expression of *PsbA*, that encodes the D1 protein, was significantly higher in CB1 plants during stress compared to IR36 **(Fig. 6C, iii)**. These results suggest that efficient *de novo* synthesis of D1 in CB1, replace damaged D1 for restoring PSII reaction centre to reinitiate photosynthesis during recovery.

### DNA methylation regulates expression of ABA dependent pathway genes during chilling stress

Plant hormones, abscisic acid (ABA) and jasmonic acid (JA), play crucial roles in the cold stress response. We measured endogenous ABA and JA levels in IR36 and CB1 under stress and recovery conditions. Under normal growth conditions, endogenous ABA levels were 232.21 ng/g in IR36 and 241.06 ng/g in CB1. After 24 hours of chilling stress, ABA levels increased to 569.74 ng/g in IR36 and 660.43 ng/g in CB1. During recovery, IR36 maintained significantly higher ABA levels (361.80 ng/g) compared to control; ABA levels reached baseline during recovery in CB1 (260.67 ng/g). Interestingly, the chilling stress response in IR36 appears to be JA-dependent, as endogenous JA increased nearly two-fold during stress, whereas no significant change in JA level occured in CB1 under either condition **(Fig. 7A, i-ii)**.

**Figure 7.**
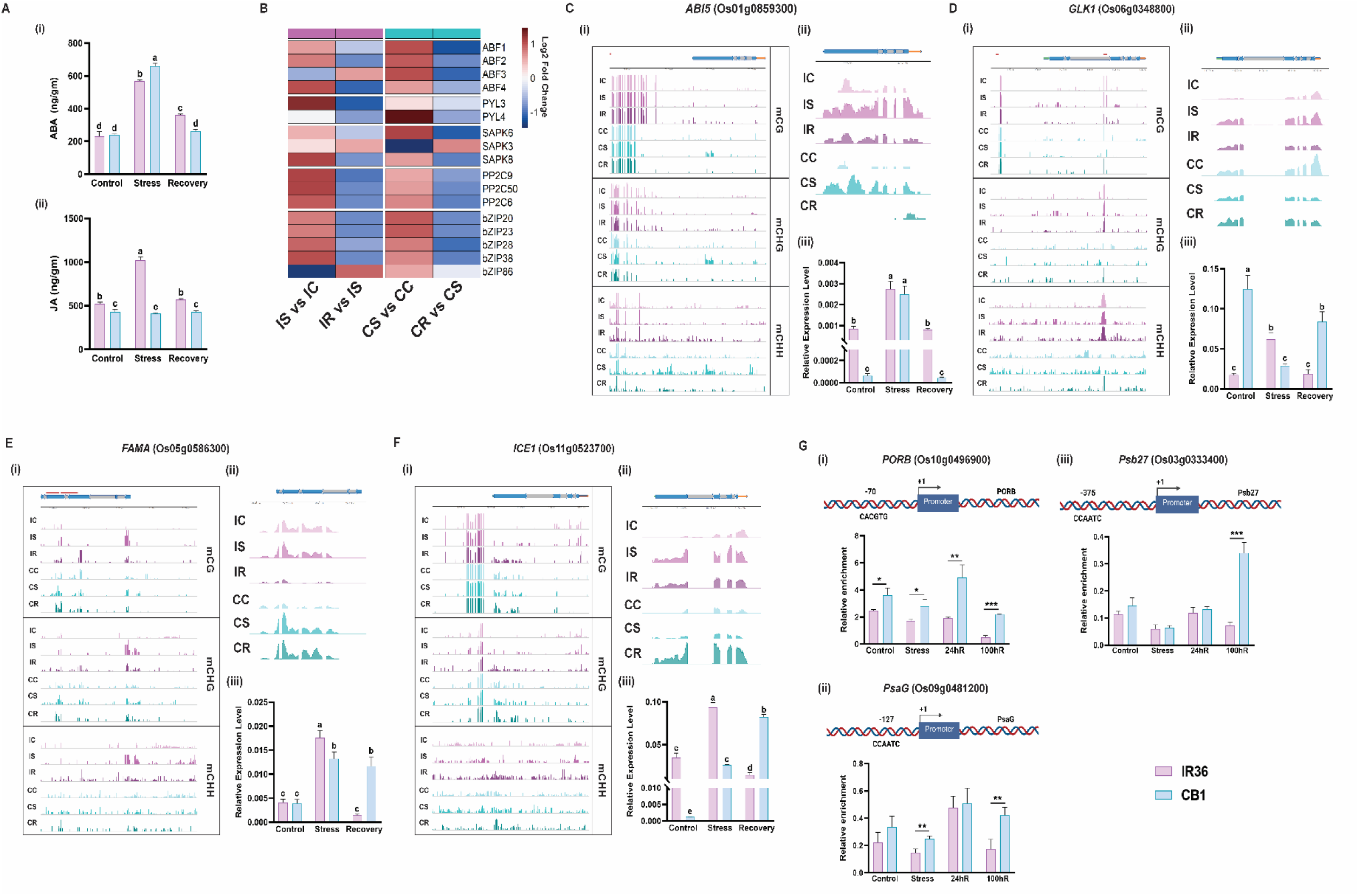
Locus specific methylation change and expression of key transcription factors during chilling stress and recovery in IR36 and CB1. **A.** Phytohormone (i) ABA and (ii) JA levels in IR36 and CB1 under control, stress, and recovery. The error bar represents mean ± S.D. from three independent experiments (*n* = 3). The significance of the data was measured by two-way ANOVA with Tukey’s post hoc test. B. Heatmap representing expression profile of DMR associated DEGs related to ABA signalling in IR36 and CB1 under stress (IS vs IC and CS vs CC) and recovery (IR vs IS and CR vs CS). The heatmap was generated using log_2_ Fold change values of respective genes**. C-F. (i)** RNA-seq coverage of respective genes in IR36 and CB1. **(ii)** RT-qPCR validation of expression of respective genes under control, stress and recovery. The error bars represent mean ± S.D. from three independent experiments (*n* = 3). The significance of the data was measured by two-way ANOVA with Tukey’s post hoc test. **(iii)** DNA methylation (CG, CHG and CHH) profiles within 2kb promoter and gene body under different conditions. IC: Control, IS: Stress, IR: Recovery for IR36 and CC: Control, CS: Stress, CR: Recovery for CB1. Red bars representing TE, Black Line (above) representing genome coordinates and horizontal bars represent gene body locations (Green: 5’UTR, Grey: Introns, Blue: Exons, Orange: 3’UTR). **G.** ChIP assay showing GLK1 occupancy at the 1 kb upstream region of **(i)** *PORB*, **(ii)** *PsaG* and **(iii)** *Psa27*. The schematic on each panels show GLK1 binding site (CACGTG/ CCAATC). The error bar represents mean ± S.D. from three independent experiments (*n* = 3). The significance of the data was measured by Two-tailed paired t-test, the level of significance was represented by \**P <* 0.05, \*\**P <* 0.01, \*\*\**P <* 0.0001.

DMR-DEG analysis identified key ABA pathway genes involved in cold stress. During chilling stress, both rice lines showed higher expression of the ABA receptor genes *PYL3* and *PYL4*; the protein kinases *SAPK6* and *SAPK8*; the protein phosphatases *PP2C6, 9, and 50*; and the downstream transcription factors *ABF1, 2, 4,* and *bZIP 20, 23, 28*, and *38*. *ABF3* and *bZIP83* were upregulated only in CB1 during stress, whereas their expression increased during recovery only in IR36. Most other genes showed basal expression during recovery **(Fig. 7B, Table S3)**. *ABI5* is an important transcription factor of core ABA signalling that acts as an ABA-dependent negative regulator of stress tolerance in rice (Zou et al., 2008). We next analysed gene expression and DNA methylation changes of *ABI5*, *DREB1A*, and *DREB1B*. Under control conditions, *ABI5* and *DREB1A* expression was significantly higher in IR36 than in CB1. Expression increased during stress and declined to the baseline during recovery in both cultivars; IR36 maintained higher expression during recovery compared to CB1 **(Fig. 7C, i-ii; Fig. S6.A, i-ii; Table S4)**. At the *ABI5* locus, methylation at promoter-region mostly remained unchanged during stress and recovery in both germplasms, except that *ABI5* promoter is less methylated in CB1 compared to IR36. For gene body, IR36 showed no methylation changes during stress or recovery. In contrast, all three methylation marks increased in CB1 during stress, which decreased during recovery **(Fig. 7C, iii & S7.A)**. At the *DREB1A* locus, promoter methylation levels showed no significant changes in either germplasm during stress or recovery. However, gene-body methylation levels, that increased significantly during stress in IR36, return to baseline during recovery. In CB1, ^m^CHG and ^m^CHH were increased in gene body during chilling stress **(Fig. S6.A, iii & S7.B)**. *DREB1B* promoter showed decreased ^m^CHH during stress in IR36 and decreased ^m^CG and ^m^CHG during recovery, whereas no significant change observed in CB1. Gene-body methylation showed significant change in methylation for all three contexts in both germplasms. **(Fig. S6.B, iii & S7.C)**. Our results demonstrate that expression key ABA-dependent and independent transcription factors are regulated through germplasm-specific DNA methylation.

### Differential expression of transcription factors regulates expression of photosynthetic genes and stomatal development during stress and recovery

Transcriptome analysis identified several key transcription factors (TFs) regulating photosynthetic gene expression were differentially regulated during stress and recovery (**Table S4**). Golden2-like (GLK), a GARP superfamily TF, coordinates chloroplast development under extreme environmental conditions (Waters *et al*., 2008, 2009). Under control conditions, *GLK1* expression was significantly higher in CB1 than in IR36. During chilling stress, *GLK1* expression decreased in CB1, but expression increased significantly during recovery. IR36 exhibited a minimal increase in *GLK1* transcripts during stress, and significant decline during recovery **(Fig. 7D, i-ii)**. Promoter methylation at *GLK1* showed increased ^m^CHG and ^m^CHH during stress in IR36 and decreased ^m^CHG during recovery. In contrast, CB1 showed no significant methylation changes during stress, whereas ^m^CHH decreased during recovery. Similarly, ^m^CG and ^m^CHG increased at the gene body in IR36 during stress, whereas CB1 maintained low ^m^CG under stress which returned to baseline during recovery **(Fig. 7D, iii & S7.D)**. Since GLK1 transcription factor mediates transcription of many key photosynthesis genes, we next investigated GLK1 occupancy at the promoter or upstream region of GLK1 targets from chlorophyll biosynthesis, and LHCs and PS II repair genes using chromatin Immunoprecipitation assay (ChIP) **(Table S5)**. The ChIP analysis indicated significant increase of GLK1 occupancy at the promoter/upstream region of *Psb27* in CB1 plants compared to IR36 during recovery. The promoter/upstream regions of the *PORB* and *PsaG* genes in CB1 plants showed relatively higher GLK1 occupancy under all conditions. Together, these data suggest that GLK1 positively regulate the expression of photosynthesis genes to re-initiate photosynthesis during recovery in CB1 plants for its growth and development **(Fig. 7G, i-iii)**.

DMR-DEG analysis also identified TFs involved in stomatal development and differentiation. *FAMA*, a bHLH TF regulating stomatal complex differentiation, was upregulated during stress in both genotypes. However, its expression decreased in IR36 during recovery, but maintained elevated levels in CB1 **(Fig. 7E, i-ii)**. SCRM/ICE1, a MYC-type bHLH TF, acts upstream of *FAMA*, regulates stomatal cell lineage initiation, proliferation, and differentiation, and servers as a master regulator of freezing tolerance (Chen and Zhang, 2026). Under control growth conditions, *ICE1* expression was significantly higher in IR36 than in CB1. Chilling stress upregulated *ICE1* in both genotypes; however, *ICE1* expression significantly declined in IR36 but continued to increase in CB1 during recovery **(Fig. 7F, i-ii)**. In *Arabidopsis*, MYB60 and MYB61 control stomatal aperture during water stress (Liang *et al*., 2005; Simeoni *et al*., 2022). *MYB61* transcript levels increased during chilling stress in both rice genotypes however, it decreased significantly during recovery in IR36. In contrast, *MYB61* expression remained high in CB1 during recovery **(Fig S6.C, i-ii)**.

Methylation dynamics at these TF loci differed between genotypes during chilling stress and recovery. For *FAMA* promoter in IR36, ^m^CHH increased during stress but return to baseline during recovery. In CB1 plants, ^m^CHH decreased during stress and recovery. Similarly, for IR36, ^m^CHG and ^m^CHH decreased at MYB61 promoter during stress and returned to baseline during recovery, whereas in CB1, ^m^CG decreased significantly during stress and all three marks increased during recovery. In the gene body, all three marks increased at *FAMA* under chilling in IR36 with no change in CB1. For *MYB61*, ^m^CHG and ^m^CHH increased in both plants. During recovery, *FAMA* maintained high ^m^CG, ^m^CHG and ^m^CHH, whereas *MYB61* maintained elevated ^m^CHG and ^m^CHH in IR36. In contrast, ^m^CHG and ^m^CHH decreased significantly at *MYB61* in CB1 **(Fig. 7E, iii; S6.C, iii; S7.E-F)**.

At the *ICE1* locus, IR36 showed no change at the promoter, whereas ^m^CHH increased during stress in CB1 and decreased during recovery. In the gene body, ^m^CG decreased in IR36, which returned to baseline during recovery, while CB1 showed a transient ^m^CHH increase during stress that returned to baseline during recovery **(Fig. 7F, iii; S7.G)**. Collectively, these results suggest that differential methylation dynamics regulate the distinct expression profiles of key stomatal transcription factors in these rice germplasms during stress and recovery.

## DISCUSSION

### CB1 indica rice exhibits tolerance against chilling temperature

Being sessile, plants are extremely vulnerable to adverse environment that affects its growth, survival and yield. Stress resilience not only depends upon stress perception but also its efficiency to recover photosynthesis, ROS homeostasis and metabolism. Although studies have deepened our knowledge on plant stress response networks, cellular physiology and molecular networks that govern plant stress recovery were less explored. In this study, we have investigated post-stress recovery in two *indica* rice: high yielding IR36 and low yielding CB1. Our results indicated that CB1 plants had significant tolerance against extreme chilling treatment, with 90% seedlings surviving after 72 hours of stress treatment, compared to 10-20% survival rate for IR36.

### CB1 plants maintain closed stomata for improving water balance during stress recovery

Regulation of stomatal aperture is very critical for controlling water loss and CO_2_ uptake for photosynthesis under water limiting conditions (Bertolino et al., 2019; Jon, 2024). Under normal conditions, stomata were smaller in CB1 compared to IR36. During chilling stress and recovery, CB1 maintained closed or partially open stomata, preserving 80% leaf relative water content (RWC); however, stomata were initially closed during stress in IR36 but remained fully open during recovery. Although fully open stomata may promote photosynthesis, it can cause a severe decrease of relative water content (RWC) in IR36 during stress recovery phase. Therefore, maintaining closed stomata may be considered as “tolerance trade-off” where high water potential is maintained during recovery but compromising photosynthesis by limiting CO_2_ intake (Henry et al., 2019). CB1 plants may utilize this unique mechanism to minimize transpiration during recovery and providing conditions for reducing oxidative damages of the guard cells.

Environmental stress often can cause ultrastructural damage of the stomata, leading to dysfunction. Hence, reprogramming of stomatal development during recovery is essential to replace damaged stomata for resuming photosynthesis and development of new leaf tissues. Stomatal development is orchestrated by three bHLH transcription factors, FAMA, MUTE and SPCH (Lampard and Bergmann, 2007). ICE1/SCRM interacts with these bHLH TFs to form heterodimers to specify their action during meristemoid differentiation, guard mother cell and guard cell formation (Kanaoka et al., 2008). Our results indicated higher expression of *FAMA* during stress recovery in CB1 plants compared to IR36. Similarly, CB1 plants also maintained high *ICE1* expression, indicating that ICE1-FAMA/MUTE/SPCH pathway was active for replacing damaged stomata after chilling stress.

### Efficient transcriptional regulation of photosynthesis during post-stress recovery in cold tolerant CB1

Comparative transcriptome between the two rice germplasms have indicated enrichment of GO terms related to photosynthesis in tolerant variety. GO terms like “pigment biosynthesis process”, “photosynthesis light harvesting photosystem I”, “photosynthesis light harvesting”, “chloroplast organization”, “chlorophyll biosynthetic process”, were uniquely enriched during recovery phase in CB1. Previous studies have shown that light and dark reactions of photosynthesis are highly challenged by abiotic factors that limit many physiological and biochemical processes (Muhammad et al., 2020). Therefore, it is assumed that plants that resumes their photosynthetic performance proficiently can recover more efficiently from environmental stress for healthy growth and development.

Chilling temperature has negative impact on chlorophyll and carotenoid content, electron transport through PS I and II, stomatal conductance and carbohydrate metabolism (Kayess et al., 2020; Seydel et al., 2022; Sonoike, 1998). In cold tolerant CB1 plants, the chlorophyll content does not change significantly during stress or recovery, compared to IR36. Further, higher expression of chlorophyll degradation genes was observed during recovery phase in IR36 plants. Studies have indicated that four chlorophyll degradation genes, *NYC1*, *NYC3*, *SGR* and *RCCR* are involved during leaf senescence and stress response, of which NYC1 and SGR are regulated by ABA. *NYC1* encodes chlorophyll b reductase and is involved in degradation of chlorophyll b and light-harvesting complex II (Sato et al., 2009). Similarly, SGR regulates disassembling of LHCII in the thylakoid membranes and its overexpression resulted in oxidative stress, leading to cell death of rice leaves at seedling stage (Jiang et al., 2011; Jiao et al., 2020; Yang et al., 2020). The low chlorophyll content of IR36 directly effects photosynthesis activity of the plant, as maximum PSII quantum efficiency declines significantly during stress and recovery, compared to CB1. Further, chilling stress severely affects the expression of photosystem core component genes in both plants; however, expression of majority of photosystem core component and LHC genes were efficiently revived in CB1 plants after stress removal. Previous studies have indicated that LHC proteins are tightly regulated by environmental stress, particularly during cold stress response (Deng et al., 2014; Meng et al., 2025). Also, overexpression of LHC genes enhanced cold tolerance of tomato plants by regulating stomatal response, ROS homeostasis, enhancing chlorophyll content and photochemical efficiency of PSII (Deng *et al*., 2014).

Extreme environmental conditions enhance photoinhibition of PSII, a process that balances the rate of photodamage vs the rate of PSII repair (Didaran et al., 2024). Photodamage of PSII involves oxidative damage and proteolytic cleavage of reaction center-binding protein, D1 that disassembles PSII-LHCII super-complex (Kale et al., 2017). The higher turnover rate of D1 promotes replacement of degraded D1 with newly synthesized protein (Inagaki, 2022). Studies have indicated that chilling stress in plants inhibits PSII repair via ROS-dependent inhibition of *de novo* D1 protein synthesis (Kale *et al*., 2017; Yang. et al., 2017). Our results indicated no significant change in in cold tolerant CB1 plants; however, D1 level decreased during stress and recovery in cold sensitive IR36. Moreover, the expression of *PsbA*, that encodes D1, increased many folds in CB1 plants during stress conditions compared to IR36.

PSII extrinsic proteins, such as PsbO, PsbP, PsbQ, and PsbR, play vital roles in protecting PSII from photodamage and stabilizing the oxygen-evolving complex (OEC) (Suorsa. and Ar, 2007). However, abiotic stressors affect these proteins (Sasi et al., 2018). Transcriptomic data revealed that most PSII extrinsic proteins were upregulated significantly during recovery in cold-tolerant CB1 plants, indicating efficient PSII damage repair to restore PSII–LHCII function. Collectively, our results suggest rapid recovery of photosynthetic apparatus of CB1 restore PSII function and oxygen-evolving capability for growth and development of the plant post stress.

### ROS scavenging system is highly active in cold tolerant CB1 plants

Chloroplast, being the major sources of ROS production (Li and Kim, 2022) maintain the equilibrium between ROS production at PSII, PSI or ETC and ROS scavenging through collective action of antioxidants, SOD, APX–glutathione cycle or thioredoxin system (Asada, 2006). Cold stress induced photoinhibition causes overloading of electron transport chain, leading to overproduction of reactive oxygen species (Wei et al., 2022). These ROS molecules are responsible for oxidative damaging of chloroplast membrane, chlorophyll destruction, inactivation of Calvin cycle enzymes, damage of Fe^2+^-containing enzymes, D1/D2 proteins and Mn-clusters in PSII (Dietz et al., 2016; Kale *et al*., 2017; Kim, 2019). Expression of some of the ROS scavenging enzymes was differentially expressed in both plants after chilling stress. Intriguingly, our results indicated that many SODs and ascorbate and glutathione peroxidases were highly expressed during recovery phases in CB1 plants, suggesting hyperactive antioxidant defence system to counter stress-induced oxidative damage, for stabilising the thylakoid membrane and photosynthetic apparatus.

### Germplasm specific DNA methylation regulates gene expression during chilling stress and post-stress recovery

Various studies have established the role of DNA methylation in regulating gene expression and silencing of transposable elements. In plants, the role of DNA methylation has been shown in plant development, response to abiotic factors and genome stability. Various studies have demonstrated the role of DNA methylation in regulating cold stress response in plants (Banerjee et al., 2017; Guo et al., 2019). 5mC DNA methylation has been shown to regulate ICE1-CBF-COR pathway during cold stress in many plant species (Liu and He, 2020; Xie et al., 2019). Expression of *Open Stomatal 1* genes that interact and phosphorylates ICE1, increases during cold stress through hypomethylation at the promoter region (Ding et al., 2015). Further DNA hypermethylation at the promoter of genes of ABA pathway and transposon silencing promotes cold tolerance in tetraploid rice (Chen et al., 2026). Further, (Song *et al*., 2025) has shown that multigeneration of cold treatment induced DNA methylation change at ACT1 promoter stably inherited in cold sensitive rice, supporting environmental directed epigenetic variation in natural selection.

Variation in DNA methylation within a similar genotype act as a key determinant for genetic diversity and natural variation. Rice has a large number of germplasms resources, that are rich in heritable epigenetic variations. DNA methylation being an important epigenetic factor contribute significantly towards phenotypic diversity, across these rice germplasms. Our analysis indicated significant differences in DNA methylation at CG, CHG and CHH sequence contexts between two germplasm IR36 and CB1, under control condition, and during stress response and its recovery. CB1 plants under normal growth conditions maintained low CG and CHG but high CHH methylation compared to IR36. Under chilling stress, methylation level changes in both the germplasms; however, cold tolerant CB1 restored its methylation during recovery whereas IR36 plants have difference compared to control. Further, when compared, there is a contrasting difference in differentially methylated regions between the two, with CB1 having fewer number of DMRs under all conditions. DEGs related to photosynthesis, LHCs, response to stress and hormone signaling were enriched with DMRs having contrasting methylation patterns (Hypo and Hyper) between two rice germplasms. Locus specific methylation changes also influence expression of ROS scavengers and transcription factors in IR36 and CB1. These findings highlight presence of unique epigenetic regulation in these two rice germplasms that regulates multiple pathways during chilling stress and post-stress recovery.

### Differential ABA-signaling regulate chilling stress response in rice germplasms

The plant hormone ABA plays a crucial role in cold stress response, although the mechanism of ABA response is very complex. Increase in endogenous ABA during cold stress has been shown to regulate various key regulators involved in hormonal cross talk, transcription activity, protein phosphorylation, and ROS homeostasis (Eremina et al., 2016; Li et al., 2022). Previous studies have indicated that cold stress increases endogenous ABA content by activating ABA biosynthesis genes (Xiang et al., 2024). In our case, we have also observed that endogenous ABA increased in IR36 and CB1 plants, with CB1 showing higher level during chilling stress. Although ABA content reverts back to baseline in CB1 after stress removal, IR36 maintained higher level compared to control. Studies have shown that JA and ABA acts synergistically through MYC2-ABI5 cascade to increase ABA accumulation during stress response (Wan et al., 2025). Also, MYC2 binds to the promoter of ABA biosynthesis gene *NCED2* and promotes its expression during cold stress response in tomato (Ding et al., 2022). Interestingly, both JA and ABA increase during chilling stress in IR36; however, JA content remained unchanged in CB1, suggesting that ABA accumulation in CB1 is JA independent.

*ABI5,* a bZIP TF, binds to ABRE element to regulate the expression of ABA dependent-stress responsive genes (Finkelstein, 2000,; Skubacz et al., 2016). Since the expression of *ABI5* is positively regulated by ABA, we have observed an increase in *ABI5* transcript during chilling stress in IR36 and CB1. Interestingly, when compared between the two germplasms, we have observed higher *ABI5* transcript in IR36 plants compared to CB1 even under normal growth conditions or post chilling stress recovery. Studies have shown that ABI5 negatively regulate photosynthesis by activating chlorophyll catabolism. Interestingly, we have observed higher expression of chlorophyll degrading proteins in IR36 during stress and recovery, that results in low chlorophyll content in these plants compared to CB1. Therefore, ABI5 dependent negative regulation on photosynthesis is predominant in IR36 during stress response.

GOLDEN2-LIKE (GLK), a GARP superfamily of MYB transcription factors has been shown as a key regulator of chloroplast development and chlorophyll biosynthesis (Waters et al., 2008). Recent studies have shown that GLK modulate transcription for plant adaptation during abiotic stress response by altering the expression of photosynthesis associated genes (Waters et al., 2009). GLK has been shown to activate the expression of ABA responsive genes *WRKY40*, *RhohB*, *COR15a* and *COR15b* genes in Arabidopsis, of which later two are involved in cold acclimation and freezing tolerance (RafiqAhmad et al., 2019). Intriguingly, WRKY 40 and GLK1 negatively regulate expression of ABI5 (Shang et al., 2010). This was evident when the expression of *ABI5* increased in *glk1gkl2* and *wrky40* mutants under ABA treatment (RafiqAhmad *et al*., 2019). Our results indicated that *GLK1* and *WRKY40* expressed significantly during stress recovery in CB1 plants compared to IR36. Thus, we hypothesized that higher expression of *GLK1* activates *WRKY40* that negatively regulates *ABI5* expression in CB1 plants. However, low expression of *GLK1* and *WRKY40* in IR36 promotes higher expression of *ABI5* during control and recovery. Further, GLK1 is considered as an ideal target for agricultural biotechnology to improve photosynthetic efficiency and stress tolerance in crop plants. Our ChIP assay indicated increased GLK1 occupancy at the promoter region of chlorophyll biosynthesis, LHCs and photosystem II repair and assembly proteins during recovery phase in CB1 plants. Taken together, these results suggest ABA dependent GLK1-ABI5 crosstalk regulates the expression of photosynthetic genes in CB1 for reviving the growth and development of rice plants after chilling stress.

*MYB61*, another R2R3 MYB tested in our study, has been shown to regulate stomatal closure through ABA-independent pathway (Liang et al., 2005). Although expression of *MYB61* increased significantly under chilling stress in both the rice varieties, CB1 maintained higher expression during recovery compared to IR36. In *Medicago truncatula*, MYB61 has been shown to activate *MtCBF4* expression during cold acclimation by inhibiting the binding of repressive MYB3 TF to the promoter of *MtCBF4.* Overexpression of MtMYB61 enhances survival rate, lowers electrolyte leakage and increase freezing tolerance (Zhang et al., 2016). Thus, higher expression of *OsMYB61* during chilling stress and recovery can be another positive regulator of cold tolerance trait in CB1 indica rice.

With the increasing population, the agricultural land for rice cultivation in tropical-subtropical area is also expanding. Thus, it has become a necessity to develop cold resilient high yielding rice varieties for high terrains like cold hilly regions where cold temperature proves to be a real challenge. High throughput omics and physiological analysis have indicated efficient photosynthetic efficiency, enhanced redox modulation and water-use efficiency of CB1 indica rice. There may be unique genetic and epigenetic networks that contribute to cold tolerance mechanism of CB1 variety. It is therefore important to investigate these networks to identify novel alleles and epi-alleles associated with stress or recovery to develop cold tolerance in *indica* rice

## MATERIALS AND METHODS

### Plant material and growth condition

Seeds of *Oryza sativa* L. ssp. *indica* cv IR36 and CB1 were surface sterilized with 0.1% (w/v) HgCl_2_, washed and germinated on water-soaked sterile gauge at 37°C in the dark for 72 hours. The germinated seedlings were transferred to sterile gauge in trays and grown in the presence of 0.25X Murashige and Skoog complete media at 28 °C ± 1 °C in 16 hours light and 8 hours dark photoperiodic cycle with 50% relative humidity and 700 lmol photons m− 2 s − 1 in a plant growth chamber.

### Chilling stress and Recovery treatments

For chilling stress, 14 days old seedlings of IR36 and CB1 were transferred to 8°C± 1°C for different time points as mentioned. For recovery, chilling stress treated seedlings were transferred to 28 °C ± 1 °C for different time periods up to 7 days. Seedlings grown and maintained at 28 °C ± 1 °C served as control. Recovery percentage was calculated on 7^th^ day after stress treatment under normal growth condition. The recovery percentage was calculated as Percentage of Recovery (%) = (No. of green seedlings alive/ Total number of seedlings used) x 100.

### Relative water content (RWC%)

To determine the relative water content, 14 days old seedlings of IR36 and CB1 after chilling stress at 8 °C± 1 °C for 24h, 48h, 72h were recovered for 7 days. Rice leaves of control, stress treated and recovery were weighed immediately (FW) after harvesting. The leaves were subsequently immersed in distilled water for 3 to 4 hours and turgid weight (TW) was measured. Next, the leaves were oven dried at 80°C for 3 to 4 hours to determine their dry weight (DW). The RWC was calculated as: RWC (%) = [(FW-DW)/(TW-DW)] x 100 (De Pascali et al., 2022). The experiments were conducted in five sets; each set contains at least 40 seedlings.

### Total ROS, MDA and Chlorophyll content

200mg leaf tissue of IR36 and CB1 of control, chilling stress and recovery samples were homogenized in 10mM Tris-Cl (pH7.2). After centrifugation, 10mM H_2_DCFDA (2′7’-Dichlorofluorescein diacetate suspended in DMSO) was added to the supernatant to a final concentration of 10µM. The fluorescence was measured at 504 nm excitation and 525 nm emission. The protein concentration was determined using the Bradford reagent. The ROS content was expressed as fluorescence units/mg of protein (Jambunathan, 2010).

Detection of ROS in plant leaves was performed using DAB (Sigma-Aldrich) following a previous method with adaptation (Daudi. and O’Brien., 2012). The de-chlorophyllized leaf samples were mounted on a slide with 50% glycerol and photographed using stereomicroscope (Leica KL300 LED) at 0.6x magnification.

MDA estimation was performed with 200mg leaf tissue of IR36 and CB1 seedlings for control, chilling stress and recovery using protocol described earlier (Huang et al., 2016).

The chlorophyll contents were measured as described previously (LICHTENTHALER. and WELLBURN, 1983) using 200mg leaf tissue of IR36 and CB1 seedlings under different conditions. Chlorophyll (a, b and total (a+b)) concentrations were measured using the following formulas: Chlorophyll a: 12.7 x A664 - 2.79 x A647, Chlorophyll b: 20.7 x A647 - 4.62 x A664, Chlorophyll (a+b): 17.90 x A647 + 8.08 x A664.

### Maximum quantum yield of photosystem II

The maximum quantum yield of photosystem II in IR36 and CB1 seedlings under control, stress and recovery was quantified with Pocket-PEA (Hansatech Instruments Ltd, UK.). The leaves of IR36 and CB1 seedlings under various conditions were dark adapted for 2 hours before the measurement. Measurements were recorded up to 1 sec using a single flash of light with photosynthetic photon flux density of 3500 μmol m^-2^ s^-1^. The minimal fluorescence level (Fo) and maximal fluorescence level (Fm) were extracted with the PEA plus software. The Fv was calculated as Fm − Fo and the maximal quantum yield of photosystem II was calculated as Fv/Fm. Readings were taken from 50 individual seedlings of each treatment.

### Stomatal traits

To examine the stomatal aperture under different conditions, transparent nail polish was applied to peel off the epidermal layer of the leaf blade of IR36 and CB1. Samples were collected from five leaves for each treatment and clear images were captured using confocal microscope. The length, width and number of the stomata were calculated using image J software. The stomatal aperture was calculated as width/ length. Total, 100 stomata from five individual leaves, each from a different seedling, were analyzed.

### RNA extraction, RT-qPCR and illumina sequencing

RNA was isolated from 14 days old rice seedlings of IR36 and CB1 from each treatment (control, stress and recovery) using Spectrum^™^ Plant Total RNA Kit (SIGMA). For RNA-seq, three biological replicates of each treatment were used for sequencing. The RNA concentration and integrity was measured through Agilent TapeStation using high sensitivity RNA ScreenTape. cDNA library preparation and sequencing were done using NovaSeq X Plus platform (Illumina Inc., CA, USA) generating 150-bp paired-end reads. For RT-qPCR, 4μg of total RNA was used for cDNA synthesis using Revert Aid Reverse Transcriptase (Thermo Scientific) and PCR was carried out using SYBR chemistry (DyNAmo ColorFlash, Thermo Scientific). *UBQ5* (Os01g0328400) gene was used as endogenous control. All reactions were performed in triplicates, and expression levels were calculated using the ΔCt method. Relative expression level for the gene of interest was calculated using 2^-ΔCT^. The primers used are enlisted in **Table S6**.

### RNA-seq analysis

For high-quality data, adapters and low-quality reads were removed from the raw reads using Ttimmomatic v0.39 (Bolger et al., 2014). The resulting paired-end clean reads were mapped to the rice genome *Oryza sativa* Japonica IRGSP1.0 (https://ftp.ensemblgenomes.ebi.ac.uk/pub/plants/release-57/fasta/oryza_sativa/dna/Oryza_sativa.IRGSP-1.0.dna.toplevel.fa.gz ) using STAR (v 2.7.10a) (Dobin. et al., 2012) with default parameters. FeatureCounts (v 2.0.3) (Liao et al., 2014) was used to determine the read numbers mapped on each gene. Differential gene expression analysis was performed using the DESeq2 R package (Love et al., 2014). To filter out the significant differentially expressed genes, threshold fold-change was set at ±1 with a p-value cut off of ≤0.05. The functional implications of the differentially expressed genes were identified using Gene Ontology (GO) enrichment analysis using ShinyGO v0.85 program (Ge et al., 2020).

### DNA extraction and enzymatic methyl sequencing (EM-Seq)

DNA was isolated from 14 days old rice seedlings of IR36 and CB1 from each treatment (control, stress and recovery) in triplicates. Total DNA from each replicate was extracted using DNeasy Plant Mini Kit (QIAGEN). Whole genome DNA methylation was performed using EM-seq and libraries were constructed following the NEBNext® Enzymatic Methyl-seq Kit (NEB # E7120). Sequencing was performed using NovaSeq X Plus platform (Illumina Inc., CA, USA) generating 150-bp paired-end reads.

### Whole-genome methylation analysis

The sequenced raw data was processed by removal of adapter, ambiguous reads and low-quality sequences to obtain high-quality reads using Trimmomatic v0.39 (Bolger *et al*., 2014). The resulting high quality paired-end reads were used for downstream work. Bismark (Kruege *et al.,* 2011) software was used to map these high-quality reads onto the reference genome (IRGSP1.0) using default parameters. After the removal of duplicate reads, methylation data were gathered for each cytosine site. Cytosines with a sequence coverage ≥ 3X were considered for downstream methylation analysis. Methylation level (ML) was calculated as sum of methylated and unmethylated read counts in each window. ML is defined as: ML (C) = reads (mC)/ (reads (mC) + reads (C)). The relative proportion of mCs in three contexts was calculated as the proportion of mCG, mCHG and mCHH on the total mC sites, respectively. Differential methylation was calculated as DMRs using DSS (Feng *et al*., 2019) software package R. Gene Ontology enrichment analysis of DMR associated DEGs was calculated using ShinyGO.

### Thylakoid membrane isolation and Western Blot analysis

500mg of dark-adapted leaf tissue of IR36 and CB1 seedlings of control, chilling stress and stress recovery were homogenized in buffer containing 0.3M sorbitol, 25mM HEPES pH 7.5), 2mM Na-EDTA, 2mM Ascorbate, 10mM NaF, 1mM PMSF. The homogenate was filtered and the suspension was pelleted out at 2500 g for 10 min. Pellets were resuspended in hypotonic buffer (25mM HEPES pH7.5, 25mM sorbitol, 5mM NaCl, 5M MgCl_2_, 5M KCl, 10mM NaF) and centrifuged at 3500 g for 10 min to pellet the thylakoid membrane. The pellet was washed twice and resuspended in the storage buffer (50mM HEPES (pH 7.5, 0.3M sorbitol, 10mM NaCl, 5mM MgCl_2_, 10mM NaF). The chlorophyll content in the final thylakoid membrane suspension was determined as above. Equal chlorophyll concentration of thylakoid membrane suspensions was separated on a 13% SDS-PAGE gel, blotted and incubated with Anti-PsbA (Agrisera, # AS05 084) followed by ECL Detection (Thermo Fisher Scientific).

### Chromatin immunoprecipitation and ChIP-qPCR

Chromatin isolated from 1gm of 14 days old leaf tissue as mentioned earlier (Dasgupta et al., 2022). The chromatin immunoprecipitation was done using anti-GLK1 polyclonal antibody (MyBioSource, #MBS9011420) and immunoprecipitated DNA was analyzed by ChIP-qPCR. The relative enrichment was quantified by normalizing the data with respect to input samples using 2^-ΔCT^ method. Three independent biological replicate samples were used for qPCR experiments. List of primers are mentioned in **Table S6.**

### Measurements of ABA and JA

For JA & ABA estimation, 14 days old seedlings of IR36 & CB1 control, stress and recovery samples were homogenized and subjected to lyophilization. 25 mg of lyophilized sample were extracted with ice-cold methanol containing internal standard. Samples were incubated for 30 minutes at 4^0^C followed by centrifugation at 12000g for 15 minutes. The supernatant was collected and subjected to vacuum dry for an hour. Dried samples were suspended in ice cold methanol and filtered with 0.22µM filter. JA & ABA content was determined by liquid chromatography–tandem mass spectrometry.

### Statistical analysis

Quantification and statistical analysis were performed using R (v4.4.1). The values shown in the figures are either means of 3 or more independent experimental replicates ± S.D. as specified in each figure. Student’s t-test and one-way and two-way analysis of variance (ANOVAs) were performed in R. Details of statistical tests are indicated in figure legend. For estimating DNA methylation differences between control, stress and recovery, methylation level per cytosine context was represented with mean ± S.E.M. and the significance of the data was measured by Wilcoxon rank-sum test with Benjamini-Hochberg (BH) correction. Differentially expressed genes (DEGs) and differentially methylated regions (DMRs) were identified under stress, by comparing stress and control samples considering control as reference and during recovery, by comparing recovery and stress samples considering stress as reference. Similarly, analysis between recovery and control was carried out using control samples as reference.

## DATA AVAILABILITY

The datasets generated during this current study are available in the NCBI Sequence Read Archive repository. For transcriptome, BioProject ID PRJNA1457795 and for DNA methylome, BioProject ID PRJNA1563969. Currently the data is not publicly available however for any information, please contact the corresponding author.

## AUTHOR CONTRIBUTIONS

Conceptualization, S.C.; Investigation, VR.; Software and data analysis, VR.; Methodology, VR, R.P and P.D.; Microscopy, R.P and V.R.; Writing-original draft, S.C and V.R.; Writing-Review & Editing, V.R., R.P., P.D., S.C. and; Funding Acquisition, S.C.

## ACKNOWLEDGEMENTS

Dr Shubho Chaudhuri and Vishal Roy would like to acknowledge Bose Institute for providing infrastructure for experiments, Madhyamgram Experimental farm facility for plant growth and propagation and data analysis. The authors sincerely thank DBT-BUILDER (2022-2027) programme: BT/INF/22/SP45088/2022 dated 17/02/2022 grant to Department of Life Sciences, Presidency University, Kolkata for Pocket-PEA system facility.

## SUPPLEMENTARY INFORMATION

Table S1. Log_10_ FPKM of DMR associated differentially expressed chlorophyll metabolism, Photosystem I &II and LHC genes.

Table S2. Log_2_ fold change of DMR associated differentially expressed antioxidant enzyme genes.

Table S3. Log_2_ fold change of DMR associated differentially expressed ABA signaling genes.

Table S4. Log_10_ FPKM of DMR associated differentially expressed transcription factors.

Table S5. Expression analysis of GLK target genes.

Table S6. List of Primers.

---

**Figure S1.**
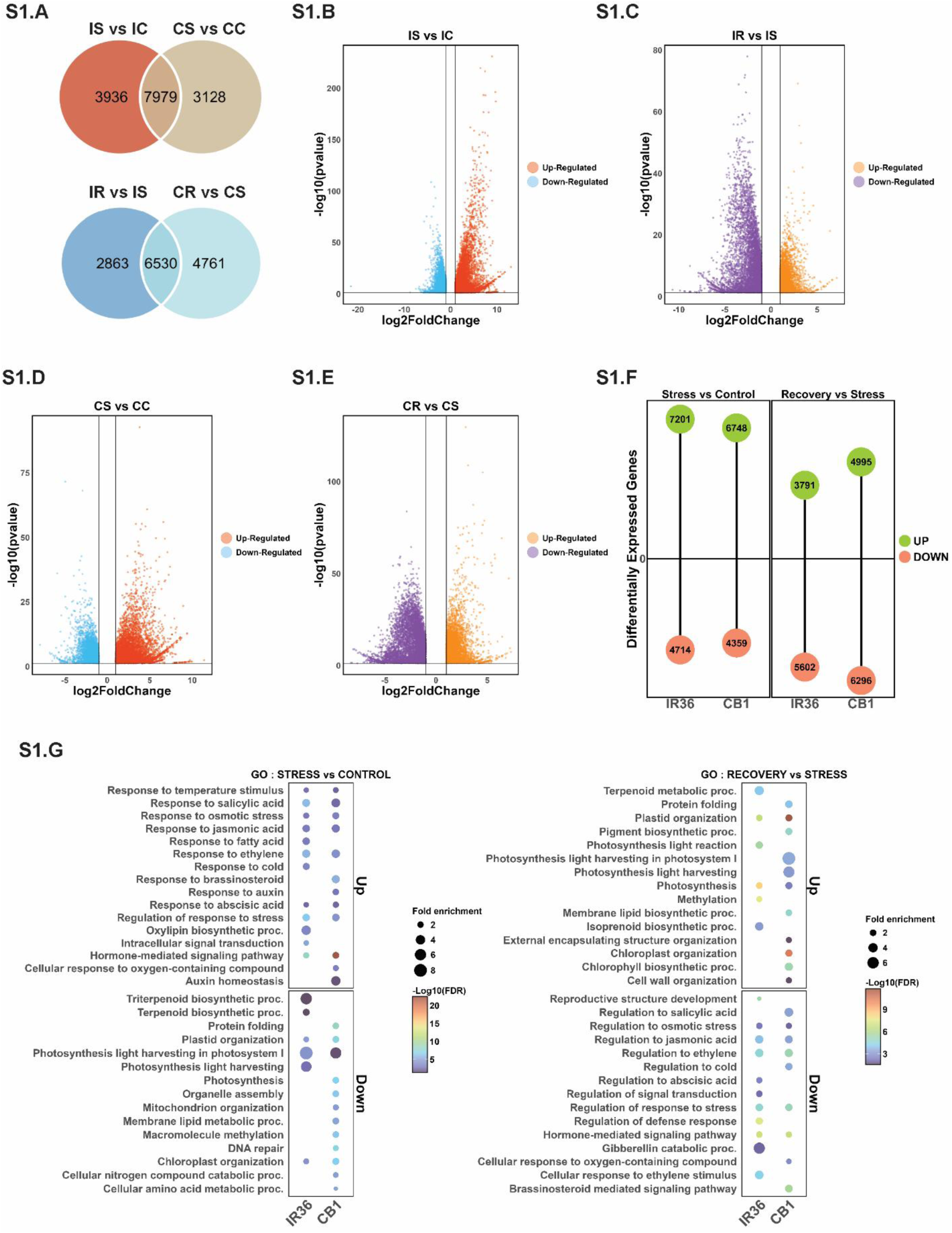
Differential Gene Expression between IR36 and CB1 under chilling stress and recovery. **A.** Venn Diagram showing the differentially expressed genes (DEGs) between IR36 and CB1 under stress (IS vs IC and CS vs CC) and recovery (IR vs IS and CR vs CS). **B-E.** Volcano plot showing up-regulated and down-regulated DEGs in IR36 and CB1 during stress and recovery. The vertical lines indicate the log2 ratio cut-off for downregulated and upregulated genes and the horizontal line represents the p-value cut off used for identifying the DEGs. **F.** Total number of upregulated and downregulated genes stress and recovery in IR36 and CB1. **G.** Dot plot showing enriched GO terms during stress and recovery.

**Figure S2.**
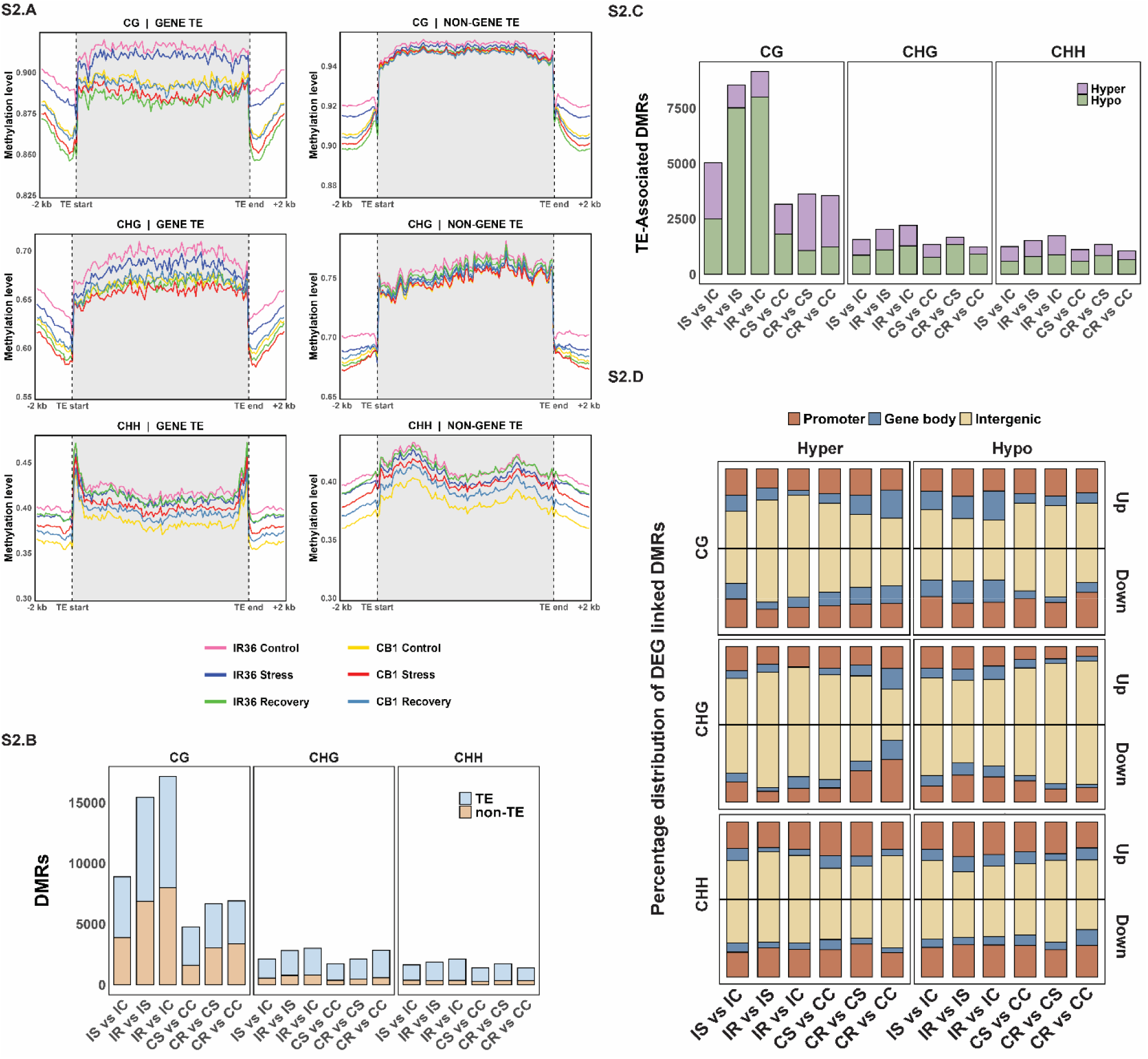
Genome wide DNA methylation at Transposon elements in IR36 and CB1. **A.** Context-specific methylation profiles of ^m^CG, ^m^CHG and ^m^CHH across genic and non-genic transposon elements (within 2 kb window) in IR36 and CB1 under control, stress and recovery. Plots aggregate signals from TE start site to TE end site in 100-bp bins. **B.** Numbers of DMRs overlapped with TE regions and non-TE regions in three sequence contexts (CG, CHG and CHH) in IR36 and CB1 **C.** Distribution of TE overlapping DMRs (Hyper and Hypo-methylated) in three sequence contexts. **D.** Distribution of TE-DMRs in different genomic regions.

**Figure S3.**
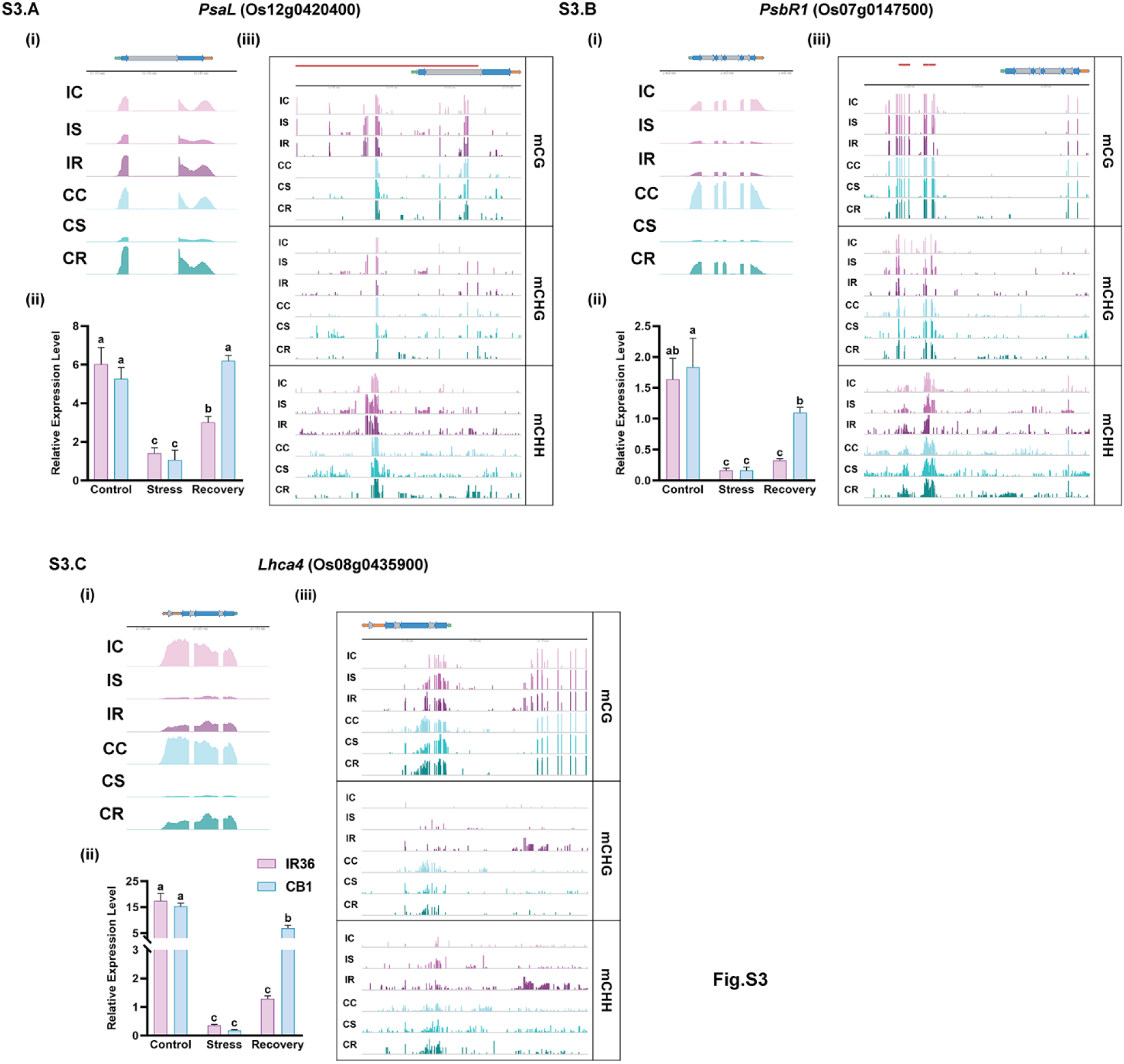
Changes in methylation and expression of photosynthesis related genes during chilling stress and recovery in IR36 and CB1. A-C. (**i**) RNA-seq coverage of respective genes in IR36 and CB1. (ii) RT-qPCR validation of expression of respective genes under control, stress and recovery. The error bars represent mean ± S.D. from three independent experiments (*n* = 3). The significance of the data was measured by two-way ANOVA with Tukey’s post hoc test. (iii) DNA methylation (CG, CHG and CHH) profiles within 2kb promoter and gene body under different conditions. IC: Control, IS: Stress, IR: Recovery for IR36 and CC: Control, CS: Stress, CR: Recovery for CB1. Red bars representing TE, Black Line (above) representing genome coordinates and horizontal bars represent gene body locations (Green: 5’UTR, Grey: Introns, Blue: Exons, Orange: 3’UTR).

**Figure S4.**
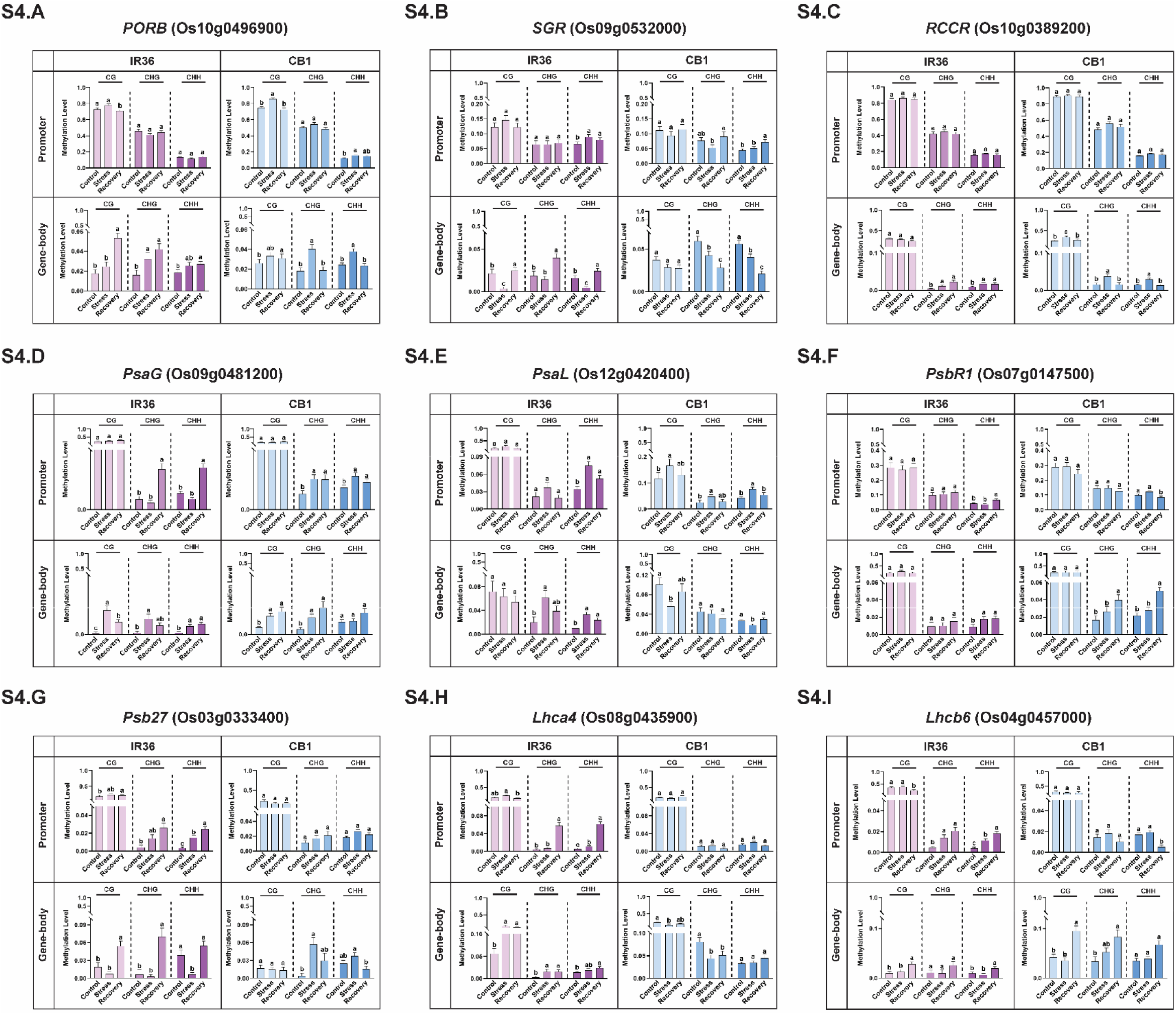
Methylation levels in photosynthesis related genes in IR36 and CB1. A-I. Bar diagram representing methylation levels of respective locus in three sequence contexts (CG, CHG and CHH) at 2kb promoter and gene body regions in IR36 and CB1 (control, stress and recovery). The error bars represent mean ± S.E.M. and the significance of the data was measured by Wilcoxon rank-sum test with Benjamini-Hochberg (BH) correction.

**Figure S5.**
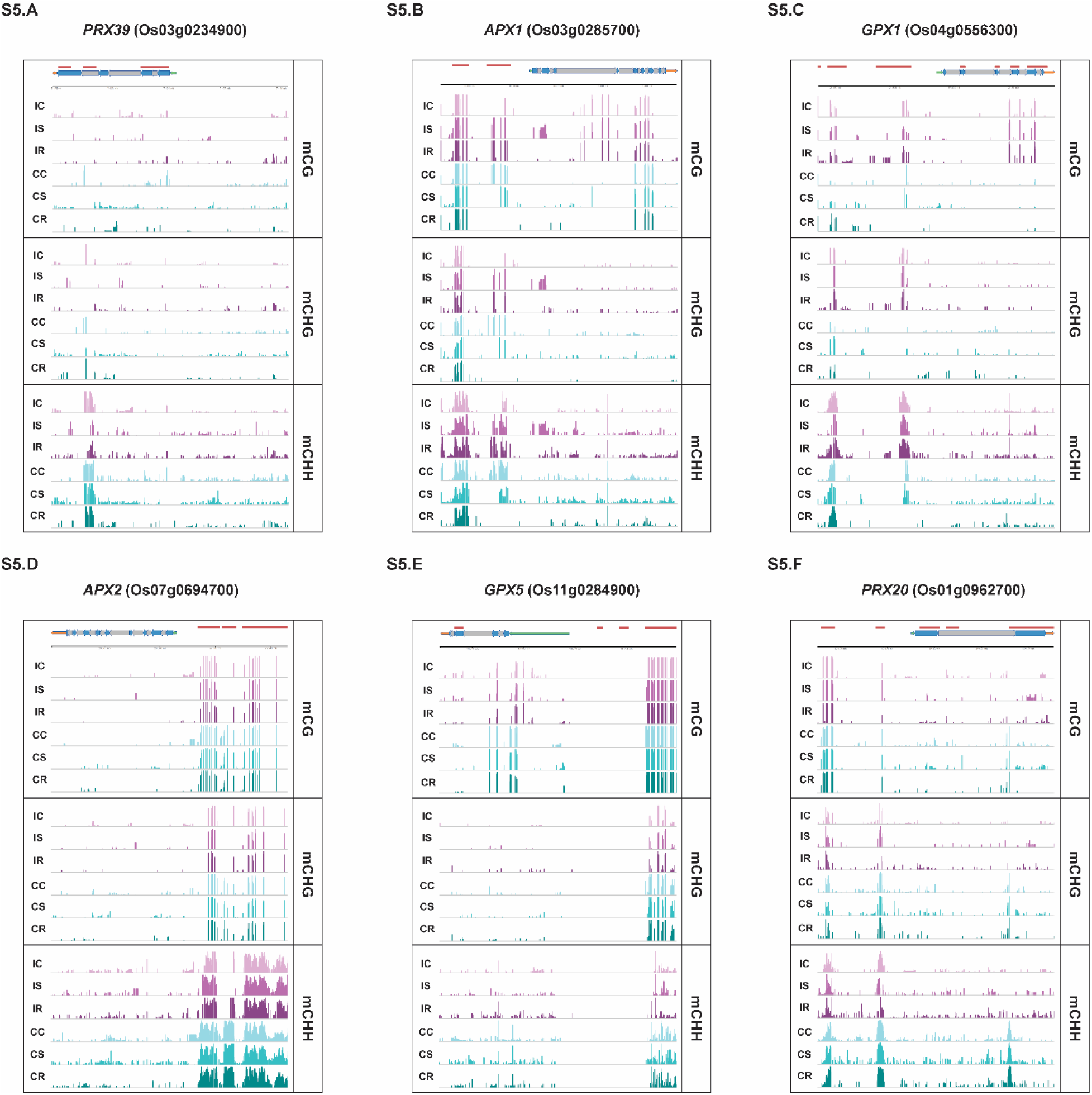
Locus specific DNA methylation changes of antioxidant genes in IR36 and CB1. A-F. Genome tracks illustrating methylation pattern of 2kb promoter and gene body in CG, CHG and CHH contexts in IR36 (Control: IC, Stress: IS, Recovery: IR) and CB1 (Control: CC, Stress: CS, Recovery: CR). Red bars (above) representing presence of TEs (Transposon Elements), Black Line (above) representing genome coordinates and horizontal bars with arrow heads (above) representing gene body locations (Green: 5’UTR, Grey: Introns, Blue: Exons, Orange: 3’UTR).

**Figure S6.**
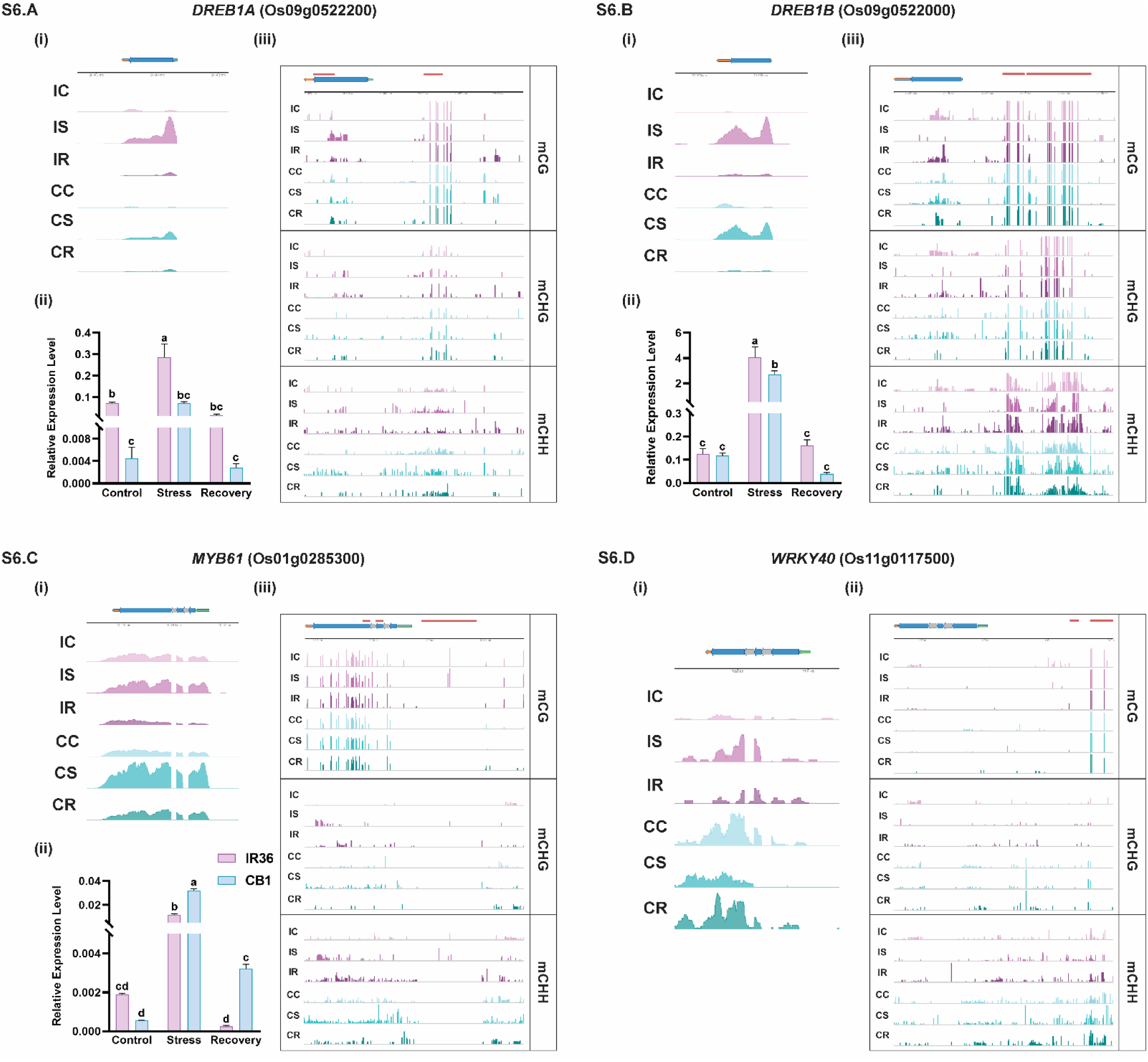
Locus specific changes of methylation and expression in transcription factors. A-D. **(i)** RNA-seq coverage of genes in IR36 and CB1. **A-C. (ii)** RT-qPCR validation of expression of respective genes under control, stress and recovery. The error bars represent mean ± S.D. from three independent experiments (*n* = 3). The significance of the data was measured by two-way ANOVA with Tukey’s post hoc test. **A-C. (iii); D. (ii)** DNA methylation (CG, CHG and CHH) profiles within 2kb promoter and gene body under different conditions. IC: Control, IS: Stress, IR: Recovery for IR36 and CC: Control, CS: Stress, CR: Recovery for CB1. Red bars representing TE, Black Line (above) representing genome coordinates and horizontal bars represent gene body locations (Green: 5’UTR, Grey: Introns, Blue: Exons, Orange: 3’UTR).

**Figure S7.**
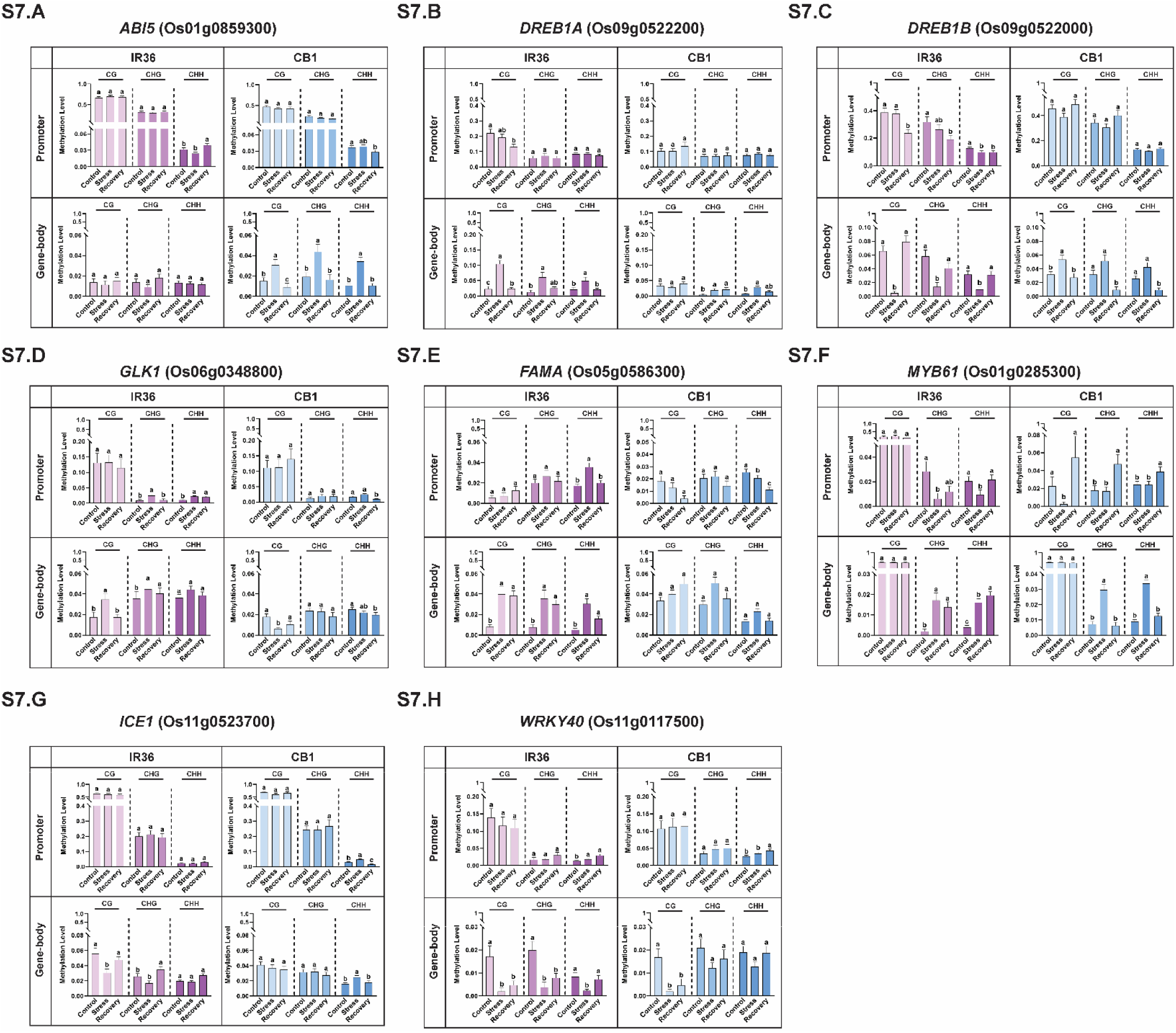
Methylation levels of transcription factors in IR36 and CB1. A-H. Bar graphs representing methylation levels of respective locus in three sequence contexts (CG, CHG and CHH) at 2kb promoter and gene body regions in IR36 and CB1 (control, stress and recovery). The error bars represent mean ± S.E.M. and the significance of the data was measured by Wilcoxon rank-sum test with Benjamini-Hochberg (BH) correction.

