## Supplementary Figures for "Differential DNA methylation and transcriptome revealed enhanced photosynthetic recovery and ROS modulation to promote cold tolerance in *Oryza sativa* L. ssp. *indica*"

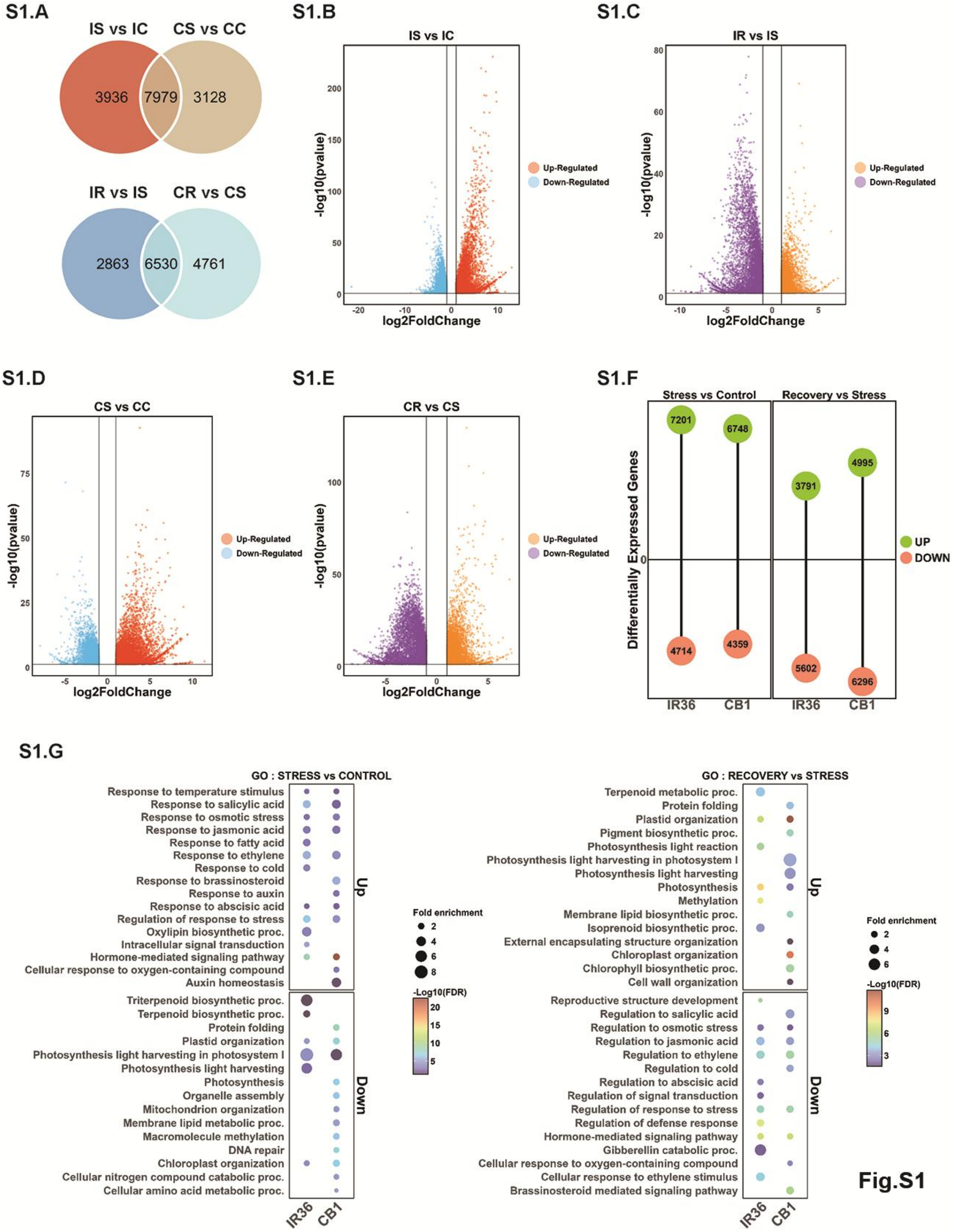
S1.A
IS vs IC
CS vs CC
3936
7979
3128
IR vs IS
CR vs CS
2863
6530
4761
S1.B
IS vs IC
-log10(pvalue)
log2FoldChange
Up-Regulated
Down-Regulated
S1.C
IR vs IS
-log10(pvalue)
log2FoldChange
Up-Regulated
Down-Regulated
S1.D
CS vs CC
-log10(pvalue)
log2FoldChange
Up-Regulated
Down-Regulated
S1.E
CR vs CS
-log10(pvalue)
log2FoldChange
Up-Regulated
Down-Regulated
S1.F
Stress vs Control
Recovery vs Stress
Differentially Expressed Genes
7201
6748
3791
4995
4714
4359
5602
6296
IR36
CB1
IR36
CB1
UP
DOWN
S1.G
GO : STRESS vs CONTROL
Response to temperature stimulus
Response to salicylic acid
Response to osmotic stress
Response to jasmonic acid
Response to fatty acid
Response to ethylene
Response to cold
Response to brassinosteroid
Response to auxin
Response to abscisic acid
Regulation of response to stress
Oxylipin biosynthetic proc.
Intracellular signal transduction
Hormone-mediated signaling pathway
Cellular response to oxygen-containing compound
Auxin homeostasis
Triterpenoid biosynthetic proc.
Terpenoid biosynthetic proc.
Protein folding
Plastid organization
Photosynthesis light harvesting in photosystem I
Photosynthesis light harvesting
Photosynthesis
Organelle assembly
Mitochondrion organization
Membrane lipid metabolic proc.
Macromolecule methylation
DNA repair
Chloroplast organization
Cellular nitrogen compound catabolic proc.
Cellular amino acid metabolic proc.
Up
Down
IR36
CB1
Fold enrichment
-Log10(FDR)
GO : RECOVERY vs STRESS
Terpenoid metabolic proc.
Protein folding
Plastid organization
Pigment biosynthetic proc.
Photosynthesis light reaction
Photosynthesis light harvesting in photosystem I
Photosynthesis light harvesting
Photosynthesis
Methylation
Membrane lipid biosynthetic proc.
Isoprenoid biosynthetic proc.
External encapsulating structure organization
Chloroplast organization
Chlorophyll biosynthetic proc.
Cell wall organization
Reproductive structure development
Regulation to salicylic acid
Regulation to osmotic stress
Regulation to jasmonic acid
Regulation to ethylene
Regulation to cold
Regulation to abscisic acid
Regulation of signal transduction
Regulation of response to stress
Regulation of defense response
Hormone-mediated signaling pathway
Gibberellin catabolic proc.
Cellular response to oxygen-containing compound
Cellular response to ethylene stimulus
Brassinosteroid mediated signaling pathway
Up
Down
IR36
CB1
Fold enrichment
-Log10(FDR)

**Fig.S1**

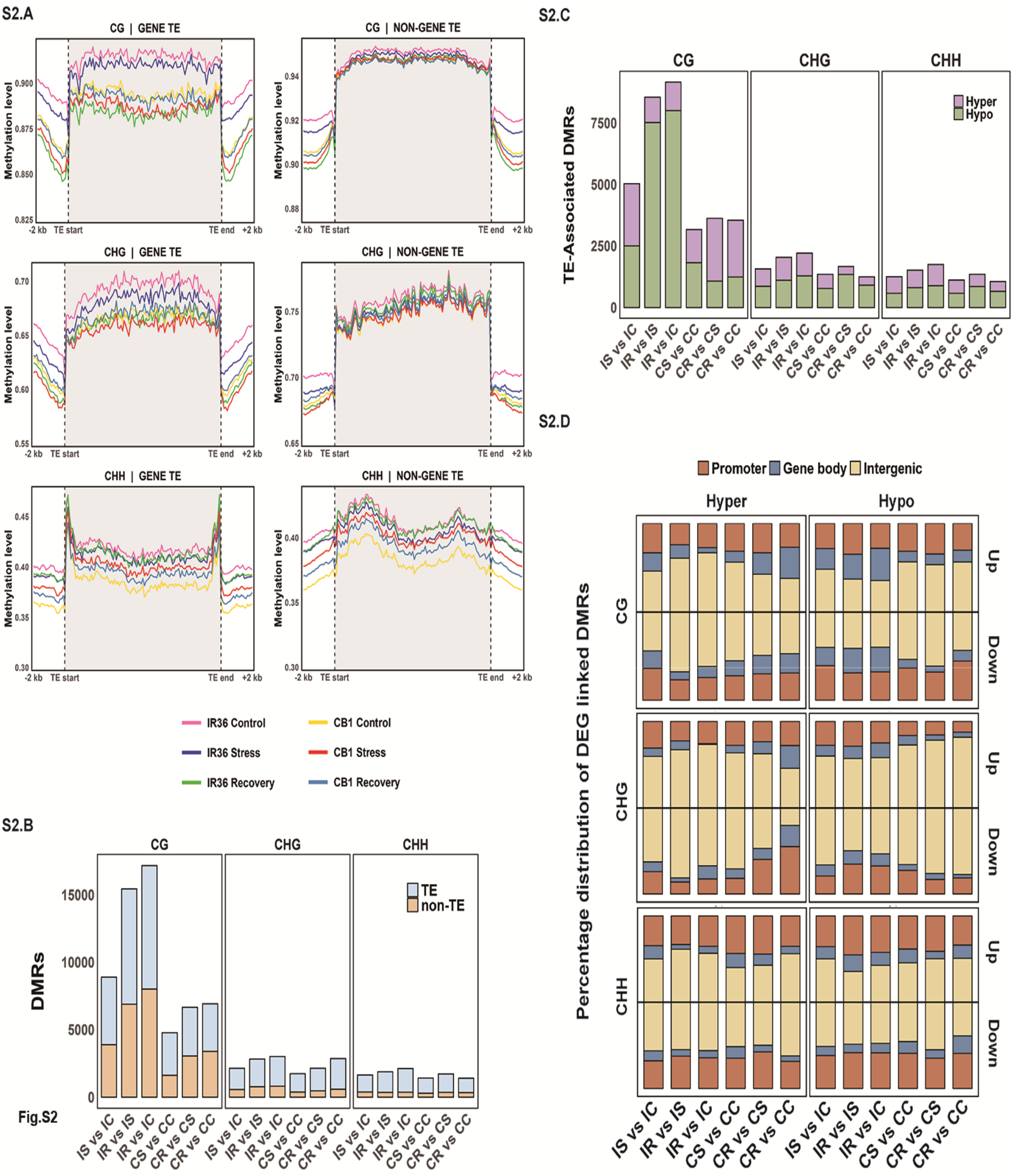
S2.A
CG | GENE TE
CG | NON-GENE TE
CHG | GENE TE
CHG | NON-GENE TE
CHH | GENE TE
CHH | NON-GENE TE
Methylation level
-2 kb
TE start
TE end
+2 kb
IR36 Control
IR36 Stress
IR36 Recovery
CB1 Control
CB1 Stress
CB1 Recovery
S2.B
CG
CHG
CHH
DMRs
TE
non-TE
IS vs IC
IR vs IS
IR vs IC
CS vs CC
CR vs CS
CR vs CC
S2.C
TE-Associated DMRs
Hyper
Hypo
S2.D
Promoter
Gene body
Intergenic
Hyper
Hypo
Up
Down
Percentage distribution of DEG linked DMRs
Fig.S2

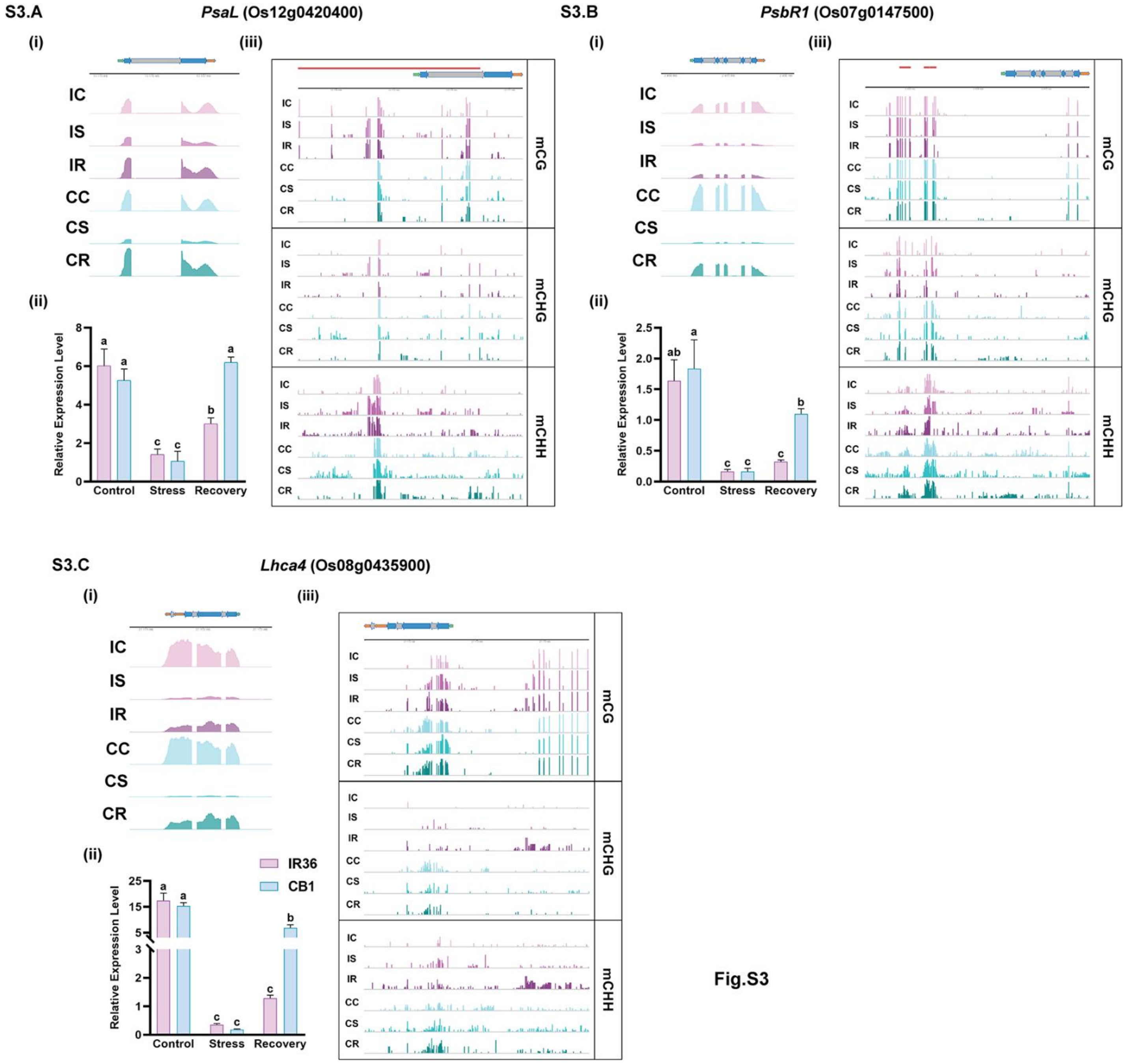

S3.A
PsaL (Os12g0420400)
(i)
(iii)
IC
IS
IR
CC
CS
CR
(ii)
Relative Expression Level
Control
Stress
Recovery
mCG
mCHG
mCHH
S3.B
PsbR1 (Os07g0147500)
S3.C
Lhca4 (Os08g0435900)
IR36
CB1


Fig.S3

Fig.S4

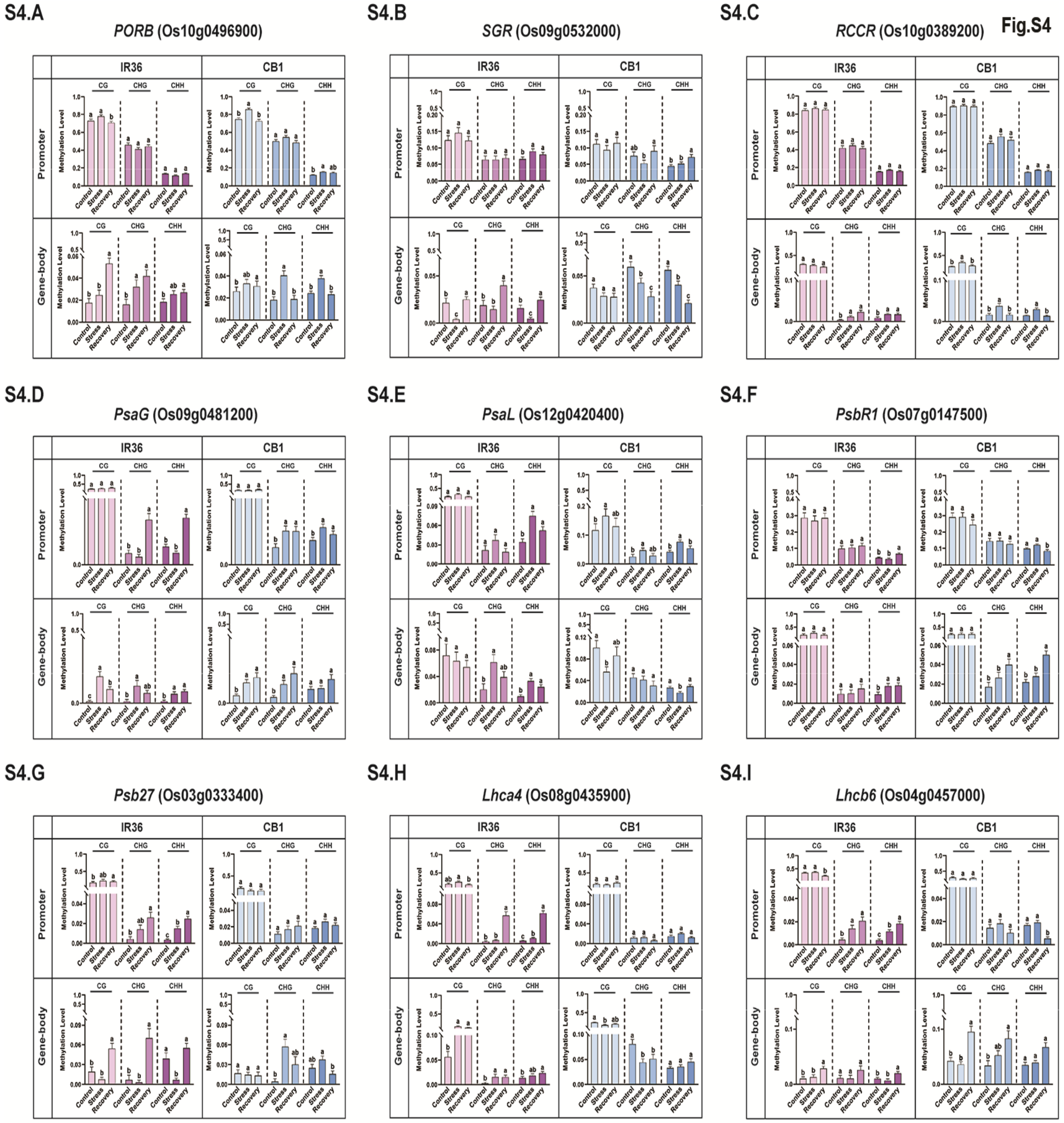
S4.A
PORB (Os10g0496900)
S4.B
SGR (Os09g0532000)
S4.C
RCCR (Os10g0389200)
S4.D
PsaG (Os09g0481200)
S4.E
PsaL (Os12g0420400)
S4.F
PsbR1 (Os07g0147500)
S4.G
Psb27 (Os03g0333400)
S4.H
Lhca4 (Os08g0435900)
S4.I
Lhcb6 (Os04g0457000)
IR36
CB1
Promoter
Gene-body
CG
CHG
CHH
Methylation Level
Control
Stress
Recovery

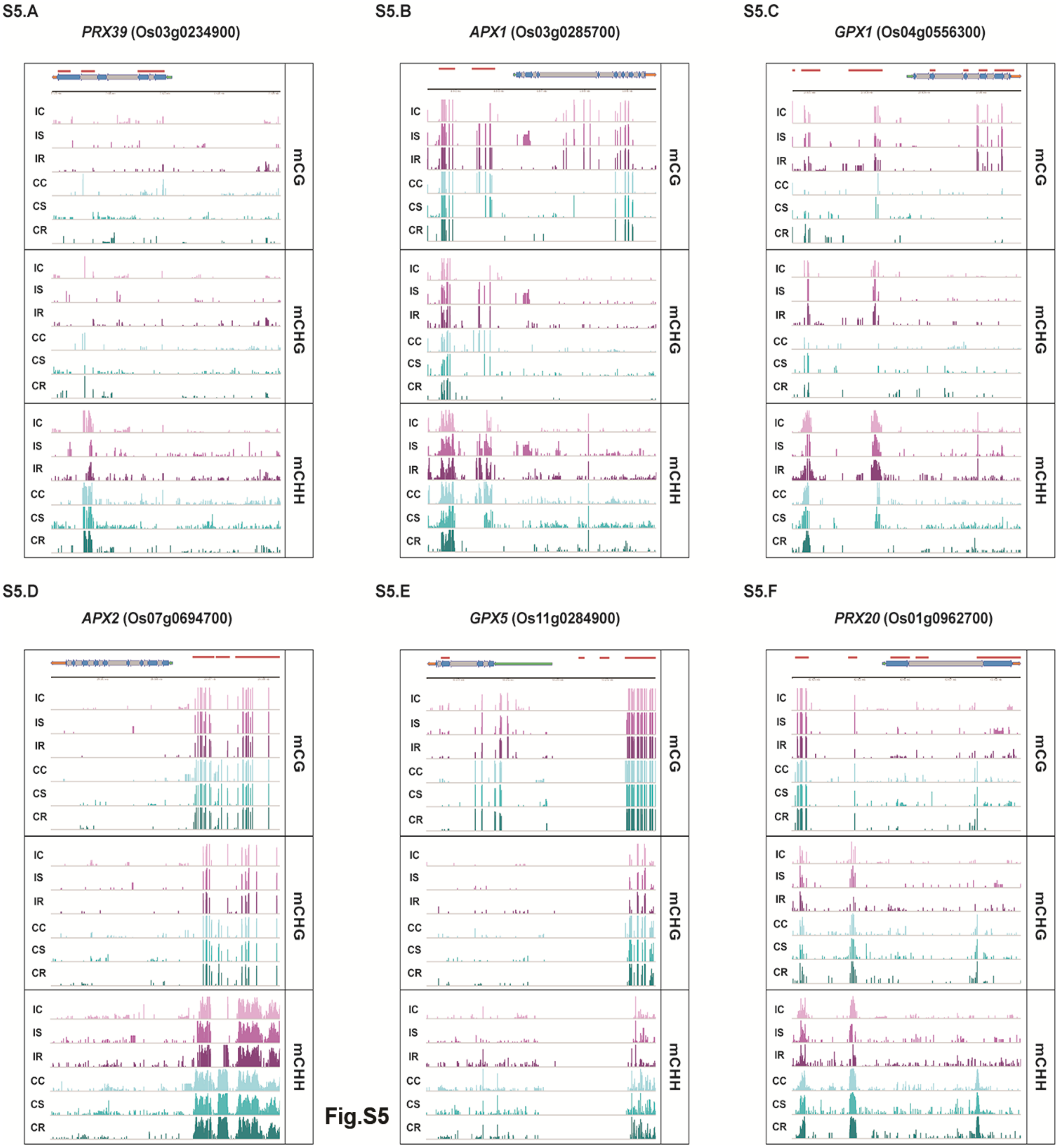
S5.A
PRX39 (Os03g0234900)
S5.B
APX1 (Os03g0285700)
S5.C
GPX1 (Os04g0556300)
S5.D
APX2 (Os07g0694700)
S5.E
GPX5 (Os11g0284900)
S5.F
PRX20 (Os01g0962700)
IC
IS
IR
CC
CS
CR
mCG
mCHG
mCHH

Fig.S5

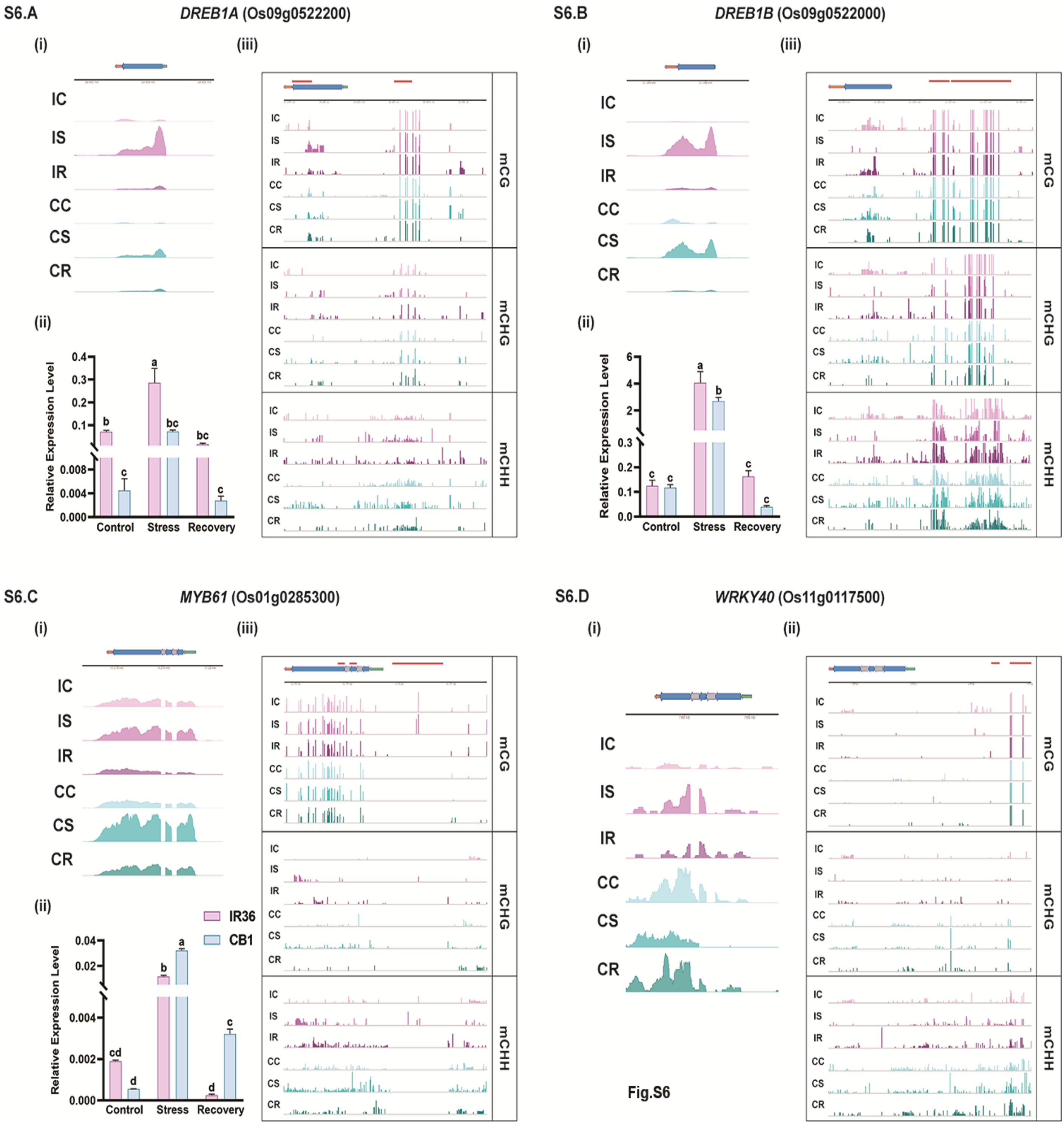
S6.A
DREB1A (Os09g0522200)
(i)
(iii)
IC
IS
IR
CC
CS
CR
(ii)
Relative Expression Level
0.4
0.3
0.2
0.1
0.008
0.004
0.000
Control
Stress
Recovery
mCG
mCHG
mCHH
S6.B
DREB1B (Os09g0522000)
(i)
(iii)
(ii)
Relative Expression Level
6
4
2
0.3
0.2
0.1
0.0
Control
Stress
Recovery
S6.C
MYB61 (Os01g0285300)
(i)
(iii)
(ii)
IR36
CB1
Relative Expression Level
0.04
0.02
0.004
0.002
0.000
Control
Stress
Recovery
S6.D
WRKY40 (Os11g0117500)
(i)
(ii)

Fig.S6

**S7.A**

*ABI5* (Os01g0859300)

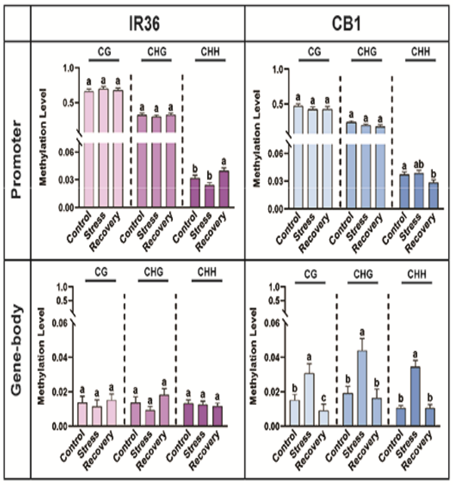


**S7.B**

*DREB1A* (Os09g0522200)

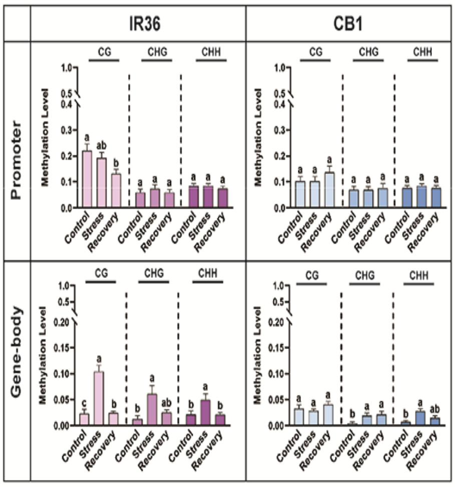


**S7.C**

*DREB1B* (Os09g0522000)

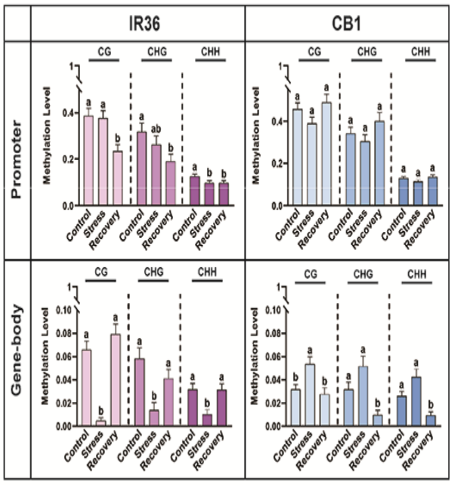


**S7.D**

*GLK1* (Os06g0348800)

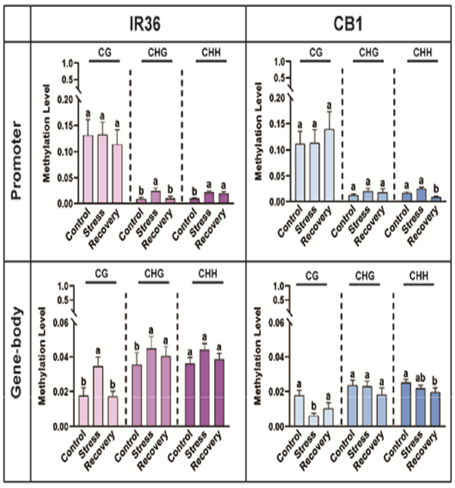


**S7.E**

*FAMA* (Os05g0586300)

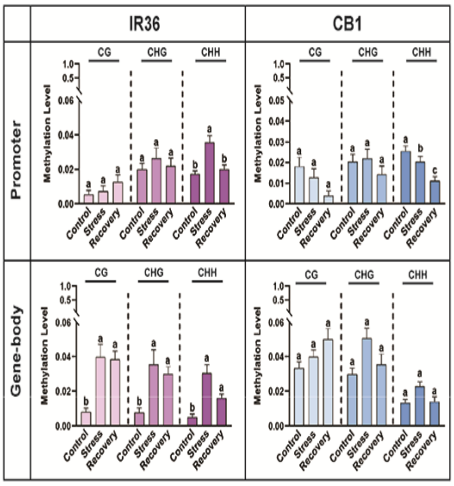


**S7.F**

*MYB61* (Os01g0285300)

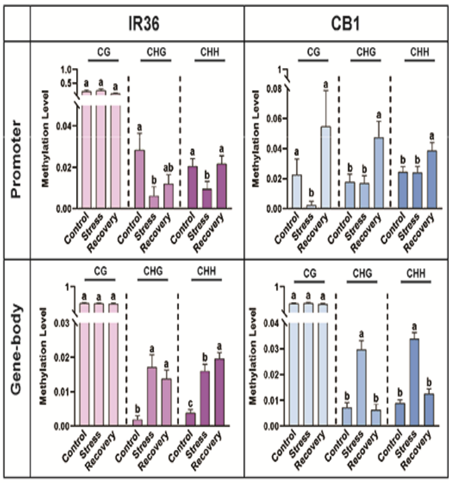


**S7.G**

*ICE1* (Os11g0523700)

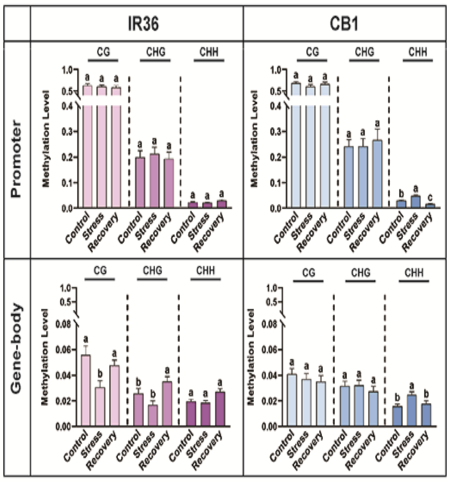


**S7.H**

*WRKY40* (Os11g0117500)

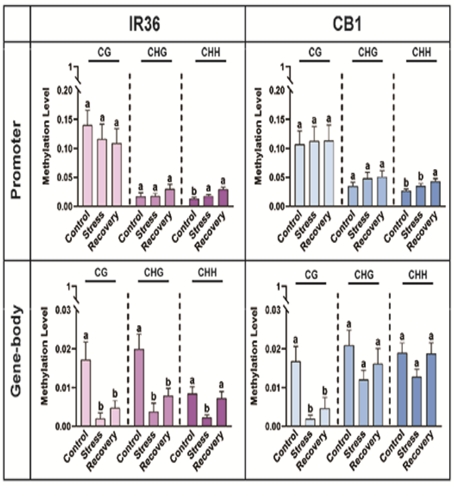


**Fig.S7**
